# Spliceosome loss in the red tide ciliate *Mesodinium rubrum* presents a symbiotic cul-de-sac

**DOI:** 10.64898/2026.07.09.736142

**Authors:** Brandon Kwee Boon Seah, Nurislam Shaikhutdinov, Wesley C. Demontigny, Erica Lasek-Nesselquist, Christiane Emmerich, Brittany N. Sprecher, Alan Kuo, Jerry Jenkins, Anna Lipzen, Kerrie Barry, Jane Grimwood, Jeremy Schmutz, Christopher Plott, Jayson Talag, Igor V. Grigoriev, John M. Archibald, Michael Lynch, Charles F. Delwiche, Holly V. Moeller, Matthew D. Johnson, Estienne C. Swart

**Author notes:** Co-first authors (Brandon Kwee Boon Seah, Nurislam Shaikhutdinov). Temasek Life Sciences Laboratory, 117604 Singapore (Brandon Kwee Boon Seah).

## Abstract

*Mesodinium rubrum*, a marine microbial eukaryote associated with some of the largest red tides on Earth, has the remarkable ability to commandeer the plastids, mitochondria and nuclei from the alga *Teleaulax amphioxeia* for photosynthesis. Here we report analyses of assemblies of *M. rubrum*’s two nuclear genomes. Unexpectedly, *M. rubrum* appears to have completely lost its spliceosomal introns, most spliceosomal molecules, and the key genes for an intron splicing-associated process, Nonsense-mediated mRNA Decay (NMD). In contrast, non-spliceosomal tRNA introns have been retained, as have thousands of intron analogs spliced out of DNA during ciliate somatic genome development (internal eliminated sequences - IESs). Intron-containing genes, especially from intron-rich species like *T. amphioxeia*, would likely be defunct if horizontally transferred to a host without a spliceosome like *M. rubrum*, and thus we propose that introns can be a roadblock to progressive endosymbiotic genomic integration.

## Introduction

Spliceosomal introns of nuclear genes are a defining eukaryotic trait inherited from the last eukaryotic common ancestor (LECA)^1^, setting them apart from bacteria and archaea. The current consensus is that the LECA was probably relatively intron-rich^2,3^. Spliceosomal introns and the spliceosome are thought to have arisen from group II self-splicing introns^4^. Multiple mechanisms can give rise to new introns^5–7^, but potentially the most rapid occurs via a subclass of non-autonomous mobile DNA elements (MITEs — Miniature Inverted-repeat Transposable Elements) known as “Introners”^8,9^ (detected in 5% of surveyed eukaryotic species^8,9^). Whatever the mechanism, once large numbers of introns became established in LECA, the spliceosome seemingly became indispensable in producing integral mRNAs.

Most reported eukaryotes with very few or no detected introns have reduced genomes (< 20 Mbp) and are typically parasites or pathogens. Eukaryotes with extreme intron reduction nonetheless still possess functional, albeit reduced, spliceosomes, e.g., *Giardia duodenalis* (12 Mbp genome, 42 introns, 4,963 protein-coding genes)^10–12^, *Encephalitozoon cuniculi* (2.9 Mbp, 37 introns, 1,997 protein-coding genes)^13,14^, *Cyanidioschyzon merolae* (a free-living red alga, 16.5 Mbp, 38 introns, 4,775 protein-coding genes)^15–17^ and *Pseudoloma neurophilia* (5.3 Mbp, 2 introns, 1,139 protein-coding genes)^18,19^ (Table 1). Kinetoplastids and prokinetoplastids have few or no canonical, cis-spliced introns, but retain a reduced spliceosome and perform trans-splicing of genes with a capped spliced-leader RNA^20–23^.

**Table 1.**
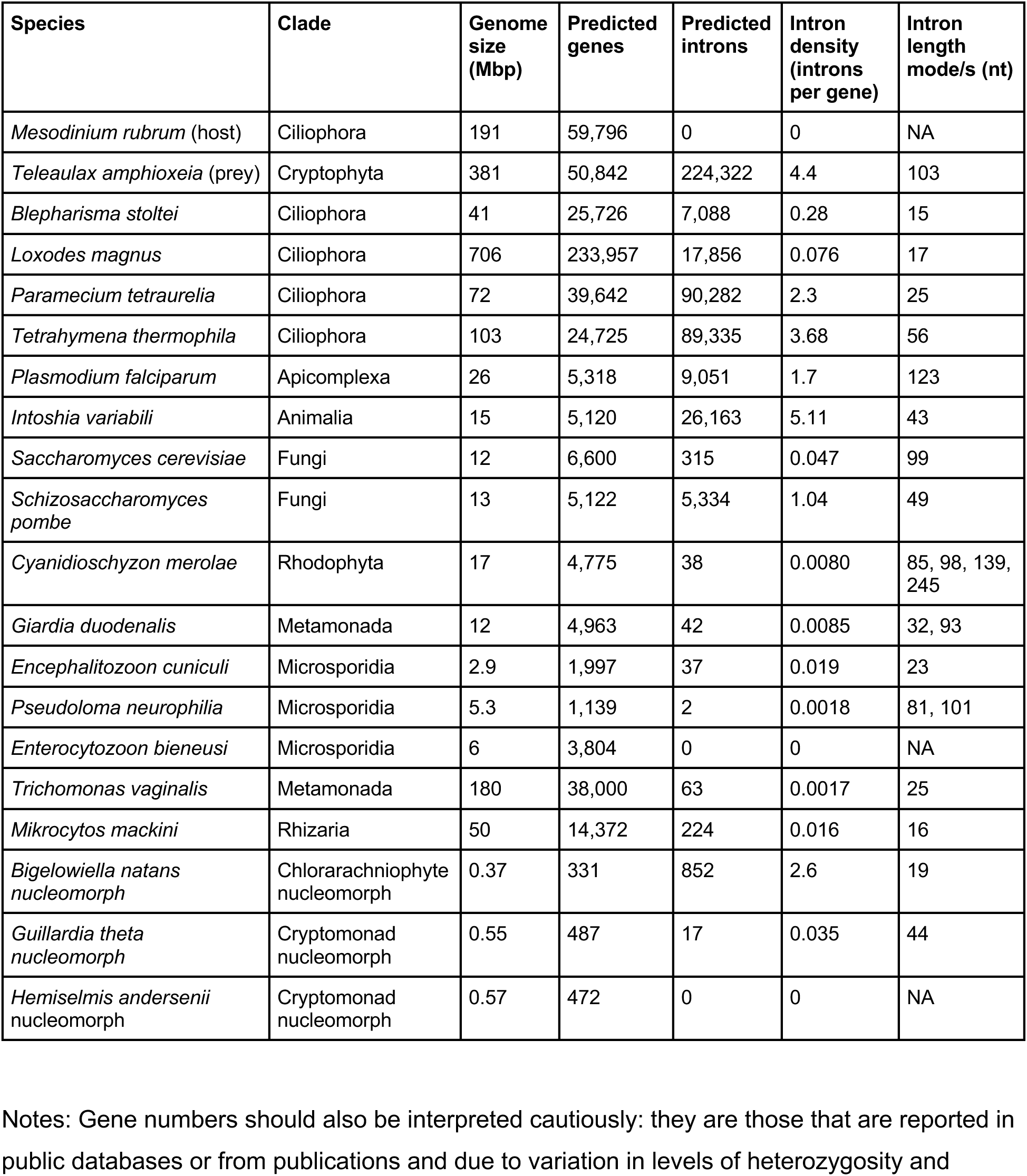
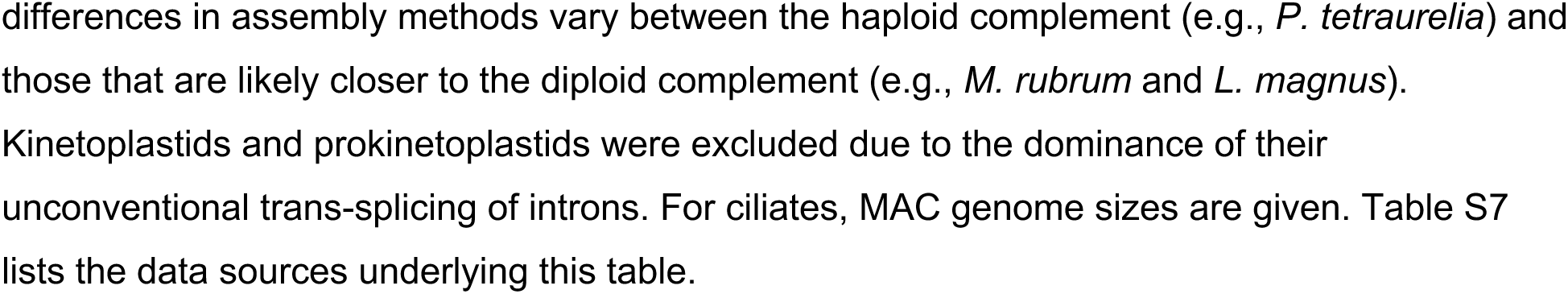
Genomic properties of organisms with few or no introns and other relevant organisms.

| Species | Clade | Genome size (Mbp) | Predicted genes | Predicted introns | Intron density (introns per gene) | Intron length mode/s (nt) |
| --- | --- | --- | --- | --- | --- | --- |
| *Mesodinium rubrum* (host) | Ciliophora | 191 | 59,796 | 0 | 0 | NA |
| *Teleaulax amphioxeia* (prey) | Cryptophyta | 381 | 50,842 | 224,322 | 4.4 | 103 |
| *Blepharisma stoltei* | Ciliophora | 41 | 25,726 | 7,088 | 0.28 | 15 |
| *Loxodes magnus* | Ciliophora | 706 | 233,957 | 17,856 | 0.076 | 17 |
| *Paramecium tetraurelia* | Ciliophora | 72 | 39,642 | 90,282 | 2.3 | 25 |
| *Tetrahymena thermophila* | Ciliophora | 103 | 24,725 | 89,335 | 3.68 | 56 |
| *Plasmodium falciparum* | Apicomplexa | 26 | 5,318 | 9,051 | 1.7 | 123 |
| *Intoshia variabilis* | Animalia | 15 | 5,120 | 26,163 | 5.11 | 43 |
| *Saccharomyces cerevisiae* | Fungi | 12 | 6,600 | 315 | 0.047 | 99 |
| *Schizosaccharomyces pombe* | Fungi | 13 | 5,122 | 5,334 | 1.04 | 49 |
| *Cyanidioschyzon merolae* | Rhodophyta | 17 | 4,775 | 38 | 0.0080 | 85, 98, 139, 245 |
| *Giardia duodenalis* | Metamonada | 12 | 4,963 | 42 | 0.0085 | 32, 93 |
| *Encephalitozoon cuniculi* | Microsporidia | 2.9 | 1,997 | 37 | 0.019 | 23 |
| *Pseudoloma neurophilia* | Microsporidia | 5.3 | 1,139 | 2 | 0.0018 | 81, 101 |
| *Enterocytozoon bieneusi* | Microsporidia | 6 | 3,804 | 0 | 0 | NA |
| *Trichomonas vaginalis* | Metamonada | 180 | 38,000 | 63 | 0.0017 | 25 |
| *Mikrocytos mackini* | Rhizaria | 50 | 14,372 | 224 | 0.016 | 16 |
| *Bigelowiella natans nucleomorph* | Chlorarachniophyte nucleomorph | 0.37 | 331 | 852 | 2.6 | 19 |
| *Guillardia theta nucleomorph* | Cryptomonad nucleomorph | 0.55 | 487 | 17 | 0.035 | 44 |
| *Hemiselmis andersenii nucleomorph* | Cryptomonad nucleomorph | 0.57 | 472 | 0 | 0 | NA |

Notes: Gene numbers should also be interpreted cautiously: they are those that are reported in public databases or from publications and due to variation in levels of heterozygosity and

Eukaryotic genomes with no spliceosomal introns at all are exceptional^19^. No introns or spliceosome components were detected in *Enterocytozoon bieneusi,* a microsporidian parasite of humans with a 6 Mbp genome (which may have experienced genome duplication)^24,25^ (Table 1). The extremely reduced, endosymbiont-derived nucleomorph genome (572 kbp) of *Hemiselmis andersenii*, a cryptophyte alga, also lacks spliceosomal introns and a spliceosome^26^). However, extreme genome reduction does not guarantee intron loss, as the nucleomorph genome of the chlorarachniophyte *Bigelowiella natans* is even smaller (373 kbp) but nevertheless has numerous (albeit short) introns (852 introns in 331 genes, modal length 19 bp)^27^. Comparisons like these, as well as additional ones considered in the discussion, challenge hypotheses that ascribe intron losses to parasitic lifestyles and genome size reduction.

The genus *Mesodinium* comprises a globally distributed, ecologically important clade of planktonic microbial eukaryotes, both freshwater and marine, that are primarily algivores and bacterivores^28^ (Figure 1A, B). The phylogenetic position of *Mesodinium* within the higher-level clade to which it belongs, ciliates, has been unclear because of its divergent small subunit rRNA (SSU rRNA) gene^29^, but phylotranscriptomic analyses currently place this genus within the litostomes, a clade that includes rumen ciliates^30^. A few *Mesodinium* species, including *M. rubrum*, sequester functional chloroplasts (kleptoplasts), mitochondria, cytoplasm, and nuclei (kleptokarya) from their preferred algal prey, rather than digesting them^31–33^ (Figure 1A). The ability of members of the *M. rubrum* species complex to harness the metabolism of their captured organelles is made possible by the functional kleptokaryon^34^, allowing these ciliates to form massive and highly productive red tides^35–38^. The relationship between *M. rubrum* and the cryptophyte alga, *T. amphioxeia*, upon which it depends, represents an intriguing form of symbiosis, with the ciliate being a kind of inverted parasitoid that destroys the alga in co-option of its organelles and photosynthetic ability^39^.

**Figure 1.**
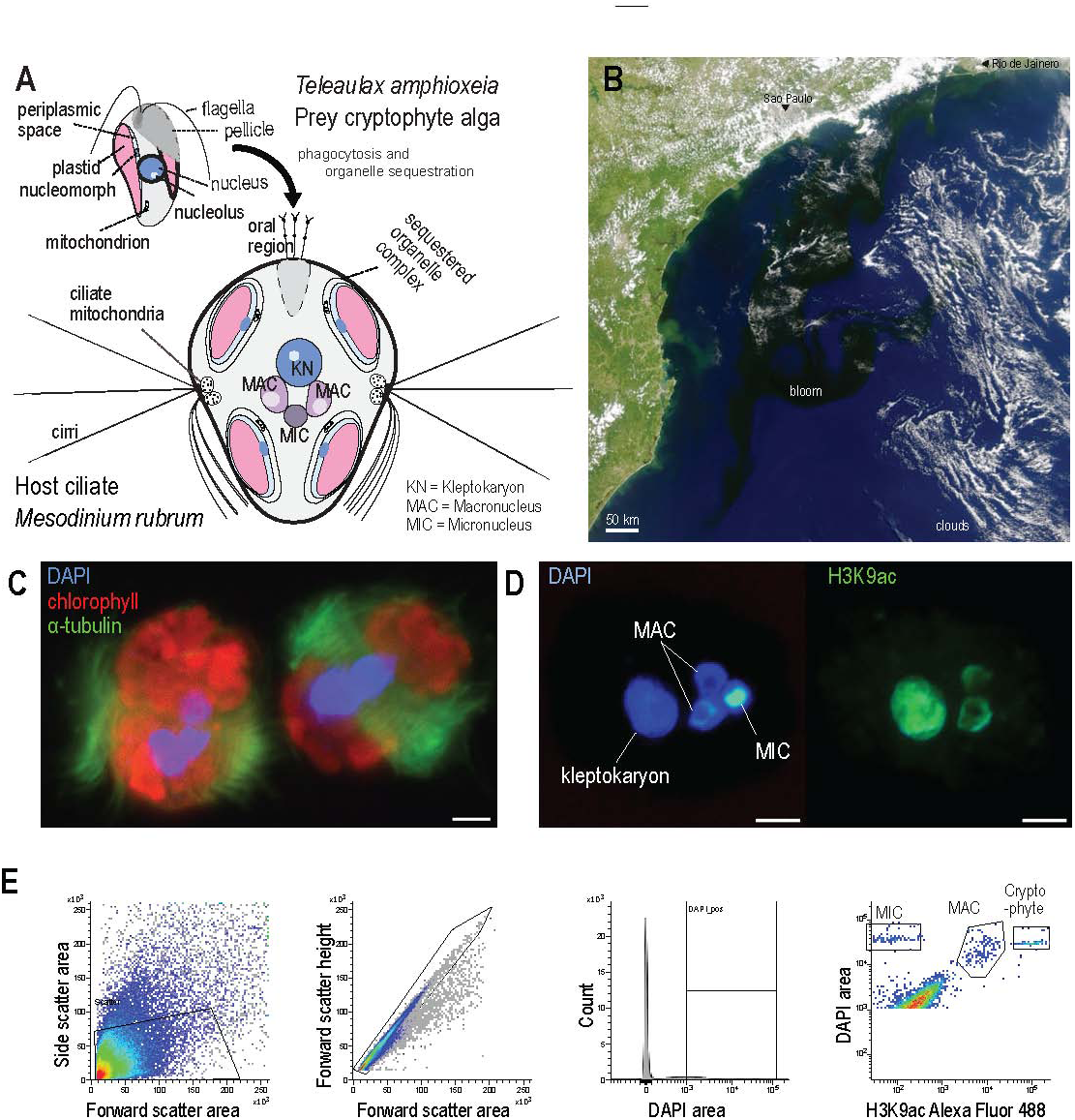
Overview of Mesodinium rubrum and purification of nuclei. (**A**) After phagocytosis of a *T. amphioxeia* cell, *M. rubrum* distributes the sequestered chloroplasts and mitochondria to the cellular periphery; captured nuclei are kept in proximity to the closely associated ciliate nuclei (MAC and MIC). Ciliate MACs develop from copies of MICs during sexual reproduction. (**B**) Satellite image of *M. rubrum* bloom off the coast of southeastern Brazil from NASA’s Aqua Modis^158^. (**C**) Fluorescence micrograph of *M. rubrum* cells; labels: alpha-tubulin immunofluorescence (green), DAPI staining of DNA (blue), chlorophyll autofluorescence (red). Scale bar: 50 µm. (**D**) *Mesodinium* and kleptokaryon nuclei stained with DAPI (blue) and by immunofluorescence for histone mark H3K9ac (green). (**E**) Gating scheme and scatter plots for flow sorting of *Mesodinium* MIC and MAC populations.

The close association between *M. rubrum and T. amphioxeia* is a stimulating case for investigation of the progression of plastid integration. Most ciliates are voracious predators that use cytostomes (“cell mouths”)^40^ to engulf (phagocytose) their microbial prey, digesting them in the resulting phagosomes^41,42^. As phagocytosis internalizes microbes, it has been suggested as a precursor to endosymbiosis and subsequent horizontal gene transfer (HGT), fitting the aphorism “you are what you eat”^43–45^. Furthermore, numerous and diverse ciliates contain algal and bacterial endosymbionts^46^. It is puzzling why fully integrated plastids (i.e., non-kleptoplasts) have not been described in ciliates, including the genus *Mesodinium*, despite hundreds of millions of years of continual phagocytosis and frequent endosymbiont possession.

We sequenced the two nuclear genomes of *Mesodinium rubrum* (strain CBJR05) to investigate the basis of its organellar capture and harnessing. To our surprise, in our initial gene predictions and mapped RNA-Seq, we were unable to find spliceosomal introns, which have been found in all published ciliate genomes. Furthermore, the absence of genes for both RNA and protein components of the spliceosomal machinery in the *M. rubrum* genome, and in the transcriptomes of *M. rubrum*, *M. pulex*, and *M. chamaeleon* suggest that spliceosomes and spliceosomal introns were likely lost in the ancestor of the genus *Mesodinium*. This stark absence of the spliceosome in these species contrasts with the variation in metabolic gene losses associated with different nutritional modes across these three *Mesodinium* species that we previously reported^39^. Loss of the ability to splice introns in this clade offers an explanation as to why cellular integration has not progressed beyond the current state.

## Results

### Purification of Mesodinium rubrum macro-and micronuclei

Like other ciliates, *Mesodinium* possesses two distinct kinds of nuclei: germline micronuclei (MIC) and somatic macronuclei (MAC) (Figure 1A). Ciliate MICs are transcriptionally inactive and, in most species, their genomes contain internally eliminated sequences (IESs), analogous to introns, but spliced out of ciliate DNA during new MAC genome development^47^. *Mesodinium* cells have small MACs similar in size to the MICs, presenting an additional challenge for genomics compared to model ciliates, such as *Tetrahymena*, *Paramecium*, *Oxytricha* and *Blepharisma,* which have large MACs containing amplified DNA. Additionally, *Mesodinium rubrum* cells retain functional nuclei (kleptokarya) from the algal prey *Teleaulax amphioxeia*. We previously investigated the metabolic capabilities of *M. rubrum*, *M. chamaeleon,* and *M. pulex* via transcriptomics^39^, with additional verification in *M. rubrum* based on the genomes we report here. Initial sequencing of *M. rubrum* whole-cell DNA yielded a complex mixture of genomes and contaminants that could not be easily separated computationally, so the nuclei had to be physically separated before sequencing.

We adapted previous flow sorting protocols^48,49^ to separate *Mesodinium* MIC, MAC, and kleptokarya. Histone H3 lysine-9 acetylation (H3K9ac), a MAC-specific histone mark in other ciliates^49–51^, was detected by immunofluorescence in *Mesodinium* MACs and kleptokarya but not MICs (Figure 1C, 1D). H3K9ac is associated with active transcription and is absent from transcriptionally silent MICs of other ciliate species like *Paramecium*^50^ and *Loxodes*^48,49^, thus *M. rubrum* MICs also appear to be transcriptionally inactive. Fluorescence signals for both H3K9ac and DAPI-stained DNA were less intense in the center of MACs, corresponding to the nucleoli^31,52^. In contrast, kleptokarya were larger and more uniformly labeled. The combination of DAPI and H3K9ac signals was therefore used to flow sort fixed, immunofluorescence-labeled nuclei from *Mesodinium* cell lysate, to obtain enriched MIC and MAC populations for sequencing (Figure 1E), verified by microscopy of the sorted nuclei.

### Organization of the Mesodinium rubrum nuclear genomes

*De novo M. rubrum* MAC and MIC draft genomes assembled by Flye^53^ were, respectively, 191 Mbp and 347 Mbp after GC% filtering to remove residual contaminants (Table S1; Figure S1). Though based on Pacific Biosciences HiFi sequencing, both genome assemblies were relatively fragmented (9,549 and 9,997 scaffolds, respectively); but MIC scaffolds (N50 45.5 kbp) were nevertheless substantially longer than those of the MAC (26.6 kbp). Assembly contiguity may have been limited by uneven coverage due to PCR amplification needed to produce sufficient DNA for genomic library preparation. Both genomes have relatively low GC base contents (MAC 31%, MIC 32%), as is typical for most published ciliate genomes^54^. This is substantially lower than the *T. amphioxeia* nuclear genome (GC = 55%).

The *M. rubrum* MAC genome assembly is larger and more repeat-rich than typical ones from model ciliate species, but there is some inflation from the true genome size due to carry-over of MIC-originating repetitive sequences, including IESs (see: “Internal eliminated sequences are enriched in the MIC genome assembly”). The presence of IESs is indicative of ciliates, not cryptophytes. A considerable portion of both assembled genomes comprises interspersed and low complexity repeats (e.g., RepeatModeller classified 104 Mbp and 240 Mbp of interspersed repeats for MAC and MIC, respectively, Table S2). Furthermore, although we chose genome assembler options to attempt to collapse alternative haplotypes, heterozygosity and partial haplotype resolution contributed to the assembly being larger than the true haploid genome size, and also to a large gene tally.

The number of protein-coding genes (<2n) predicted in the *M. rubrum* MAC (59,796) and MIC (89,847) genomes is relatively high compared to other ciliates, but not the highest (Table 1). RNA-seq mappings supported 48% versus 41% of genes in MAC and MIC assemblies respectively, with the caveat that no RNA-seq was obtained for conjugation, a ciliate life cycle phase where numerous normally quiescent genes are massively upregulated^55,56^. A noteworthy fraction of the genes in both genome assemblies is mobile-element-associated: ∼28% (MAC) and ∼23% (MIC) of proteins with domains identified are associated with mobile-element domains (Table S2 and S3). Synonymous substitution rate distributions and gene collinearity do not support the presence of a whole genome duplication (Figure S2A, B).

Predicted genes in the MIC genome assembly are shorter than in the MAC (Table S1), which may be due to a fraction of gene predictions being interrupted by IESs as well as more false positive predictions because of the high proportion of repeats (Table S2). Consistent with the latter, fewer MIC genes (20%) had identifiable Pfam domains than MAC (27%), and both were below that in predicted MAC genes of other ciliates: 45%, 49%, and 55% in *Tetrahymena thermophila*, *Oxytricha trifallax,* and *Blepharisma stoltei*, respectively. This suggests that a higher fraction of *M. rubrum* genes may have diverged beyond the domain detection capabilities of the software we used, arose as mispredictions, or originated *de novo*. The first possibility is consistent with highly divergent rRNA genes in *M. rubrum*^29^ and would also arise due to the presence of divergent mobile-element genes and pseudogenes.

*Mesodinium* belongs to the class Litostomatea^30^, which includes rumen ciliates that have obvious and abundant telomeres^57,58^. We were, however, unable to detect obvious telomeric repeats like those found in rumen or model ciliates in either of the *M. rubrum* nuclear genomes. We could neither detect homologs of the telomerase noncoding RNA (TERC) with Infernal^59^ searches, nor the corresponding telomerase protein (TERT) with TBLASTN searches using *Tetrahymena thermophila* TERT as the query (E-value < 1e-5). Exact searches of at least two successive rumen ciliate telomeric repeats, i.e., “(CCCCAAT)_2_”, and their reverse complements found zero matches in either *M. rubrum* genome assembly. Thus, it appears that both telomeres and the necessary telomere synthesis enzymes may be absent or have diverged beyond our ability to detect them with the methods we used.

Helitrons are abundant in the assembled *M. rubrum* genomes (Supplementary Results SR3; Table S2; Table S3); such mobile elements were recently shown to be inserted within degenerate telomeric repeats at chromosome ends in the *Paramecium tetraurelia* MIC genome^60^. So, it is conceivable that if *M. rubrum* has a similar germline genome organization, it could have masked the presence of telomeric repeats and decreased the assembly contiguity.

Furthermore, we observed no sign of the extremely fragmented, “nanochromosomal” genomic architecture with short telomeres observed in rumen ciliates like *Entodinium*^57,58^, although not all rumen ciliate species share this architecture^58^. The absence of short telomeres and highly fragmented MAC genomes in *M. rubrum* may be due to the absence of extensive macronuclear DNA amplification (ampliploidy; see below), which is characteristic of rumen ciliates and other ciliates with nanochromosomal architectures. Previously, telomeres, TERT, and TERC were not detected in *Loxodes magnus* nuclear genomes^49^. This ciliate species belongs to a different class (Karyorelictea), which also does not appear to have extensive DNA amplification and whose genomes are organized as long, repeat-rich, multi-gene DNA molecules^49^.

Unlike other sequenced ciliates, *M. rubrum* MACs appear to have less DNA than their MICs, and hence may lack ampliploidy. The median DAPI fluorescence of MACs was 40-80% that of MICs across different flow sorting runs (Figure 1), consistent with the relative size of the MAC to MIC genome assemblies (55%). Apart from technical error, which is also impacted by the formaldehyde fixation of the samples^61^, additional variability may be due to varying proportions of MIC contamination between runs or the presence of pre-division MACs in actively dividing cells. As the MIC and MAC assemblies have similar GC compositions, it is unlikely that the AT-preference of DAPI^62^ has affected the fluorescence-based relative DNA quantification of the *Mesodinium* nuclei. *M. rubrum*’s low MAC DNA content in relation to nuclear size and genome architecture is considered in Supplementary Discussion SD1.

*M. rubrum* has an abundance of Regulator of Chromosome Condensation 1 (RCC1) domain proteins (914 - MAC; 1234 - MIC), rivaling the most abundant mobile-element proteins (Supplementary Results SR4, Table S3). RCC1 homologs are abundant in dinoflagellates^63^; as indicated by their full name, they influence eukaryotic chromosome state, and so were suggested to contribute to dinoflagellate chromosome condensation^64^. Analysis of birefringence in *M. rubrum* cells using an LC-PolScope did in fact suggest structural ordering around the periphery of nuclei and within an internal zone, but not the conspicuous liquid crystal chromosome 3D organization found in dinoflagellates with huge genomes^65^ (Supplementary Results SR5, Figure S3). Possible roles of *M. rubrum* RCC1 proteins are considered in Supplementary Discussion SD6. Some of the most conspicuous birefringent features we observed in *M. rubrum* were ordered structures associated with their many sequestered organelle complexes (i.e., containing plastids), which may be ciliate microtubules (e.g., Figure S3C).

Although genome completeness estimates using BUSCO^66^ and a reference alveolate sequence database were well below 100% (e.g., a third of BUSCOs were not reported in the *M. rubrum* MAC genome assembly; Table S4; Data S1A), RNA-seq mapping rates to this genome were nevertheless high (90%). BUSCO statistics for *de novo* transcriptome assemblies from *M. rubrum*, *M. chamaeleon*, and *M. pulex*^39^ and the MIC genome assembly were comparable to those of the *M. rubrum* MAC genome assembly (Table S4). BUSCO completeness scores are expected to be lower than 100%, as the absent markers include genuine losses, including previously reported metabolic pathway components^39^ as well as some of the 97 spliceosomal protein orthologs sought (i.e., 11 of the missing 34 BUSCOs in the *M. rubrum* MAC genome assembly correspond to spliceosomal proteins; Table S4; see below).

### Absence of spliceosomal introns in Mesodinium rubrum

We searched for potential splice junctions by split-read RNA-Seq mapping to the *M. rubrum* MAC genome (and its prey *T. amphioxeia*), and compared these to other litostome ciliates and representative eukaryotes with few introns (*C. merolae*, *P. neurophila*, *E. cuniculi*). Split-read junctions in *M. rubrum* did not have expected ciliate intron characteristics (Figure 2; Figure S4): (i) although 481,812 potential splice junctions were found in *M. rubrum*, reflecting the high RNA-Seq coverage (mean depth 216×), most (63.3%) were supported by only one or two distinct reads; (ii) most did not have canonical GT-AG splice site motifs; and (iii) there was no obvious peak in their length distribution, which instead sloped down from the minimum length cut-off; local peaks at multiples of 3 likely represent indels due to read mapping from alternative alleles or homologs (Supplementary Results SR1). In contrast, other litostome ciliates had splice junctions that were bounded by the GT-AG motif, and which had one or more prominent peaks in the length distribution within the expected size range for ciliate introns.

**Figure 2.**
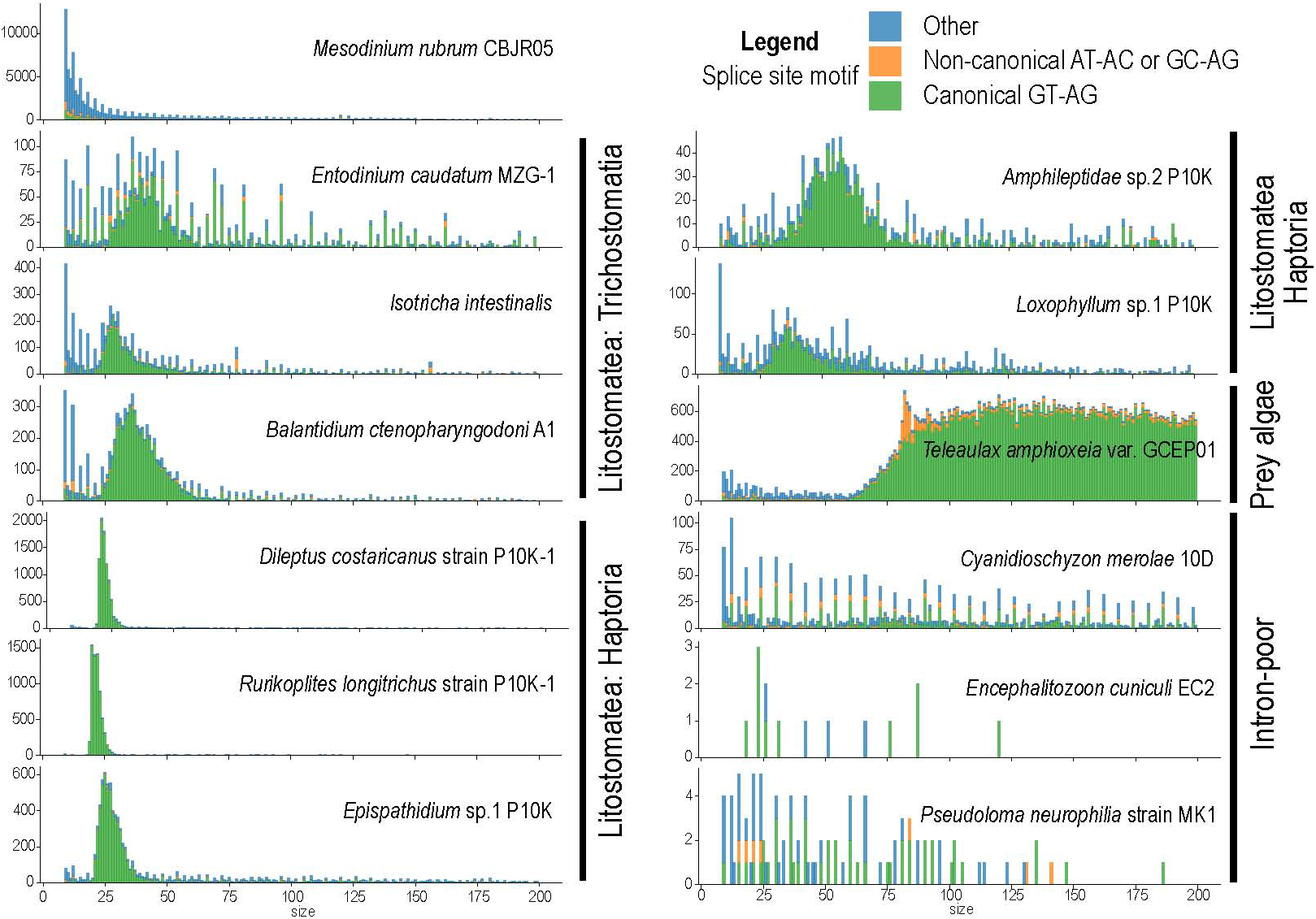
Absence of introns in mapped Mesodinium rubrum *RNA-Seq.* Length distributions of splice junctions (≤200 bp shown) supported by >2 reliable reads predicted by Portcullis from RNA-Seq mappings to the somatic genome assemblies of *M. rubrum*, other litostome ciliates, *T. amphioxeia*, and intron-poor eukaryotes. A lower intron length cut-off of 9 bp accommodates the shortest ciliate introns. Stacked bars colored by splice sites: canonical GT-AG (green), non-canonical AT-AC or GC-AG (orange), other (blue).

With RNA-Seq mapping alone, we could not rule out the possibility that *M. rubrum* may simply have very few introns that were not detectable against the background of other alignment gaps that cause split mappings, as observed with the intron-poor eukaryotes *C. merolae* and *E. cuniculi*. We therefore next searched for spliceosomal complex molecules.

### Absence of spliceosomal components in Mesodinium spp

We first searched for spliceosomal RNAs in the *M. rubrum* MAC and MIC genome assemblies using Infernal^59^. Unlike the *Tetrahymena* MAC genome, where known U1, U2, U4, U5 and U6 spliceosomal RNA genes were retrieved with minimum E-values ranging from 4.8×10^-33^ to 3.9×10^-17^, we found no compelling spliceosomal RNA gene candidates in *M. rubrum*, i.e., minimum E-values ranged from 8.3×10^-5^ to 2.5×10^-4^ for the MAC (Table S5; Data S1B to D).

Next, we searched for orthologs of known spliceosomal proteins encoded by the *M. rubrum* MAC genome, and in *de novo* assembled transcriptomes of *M. pulex*, *M. chamaeleon*, and another *M. rubrum* strain^39^ using BLAST and OrthoFinder. BLASTP reciprocal best hits identified just three orthologs of known human spliceosomal proteins in *M. rubrum* and four in *M. pulex* and *M. chamaeleon*, compared to 100 orthologs in the ciliate *T. thermophila* (Data S1E). OrthoFinder identified a few additional potential orthologs in the *M. rubrum* genomes, but these were not consistently detected in other *Mesodinium* species’ transcriptomes (Figure 3; Data S1F).

**Figure 3.**
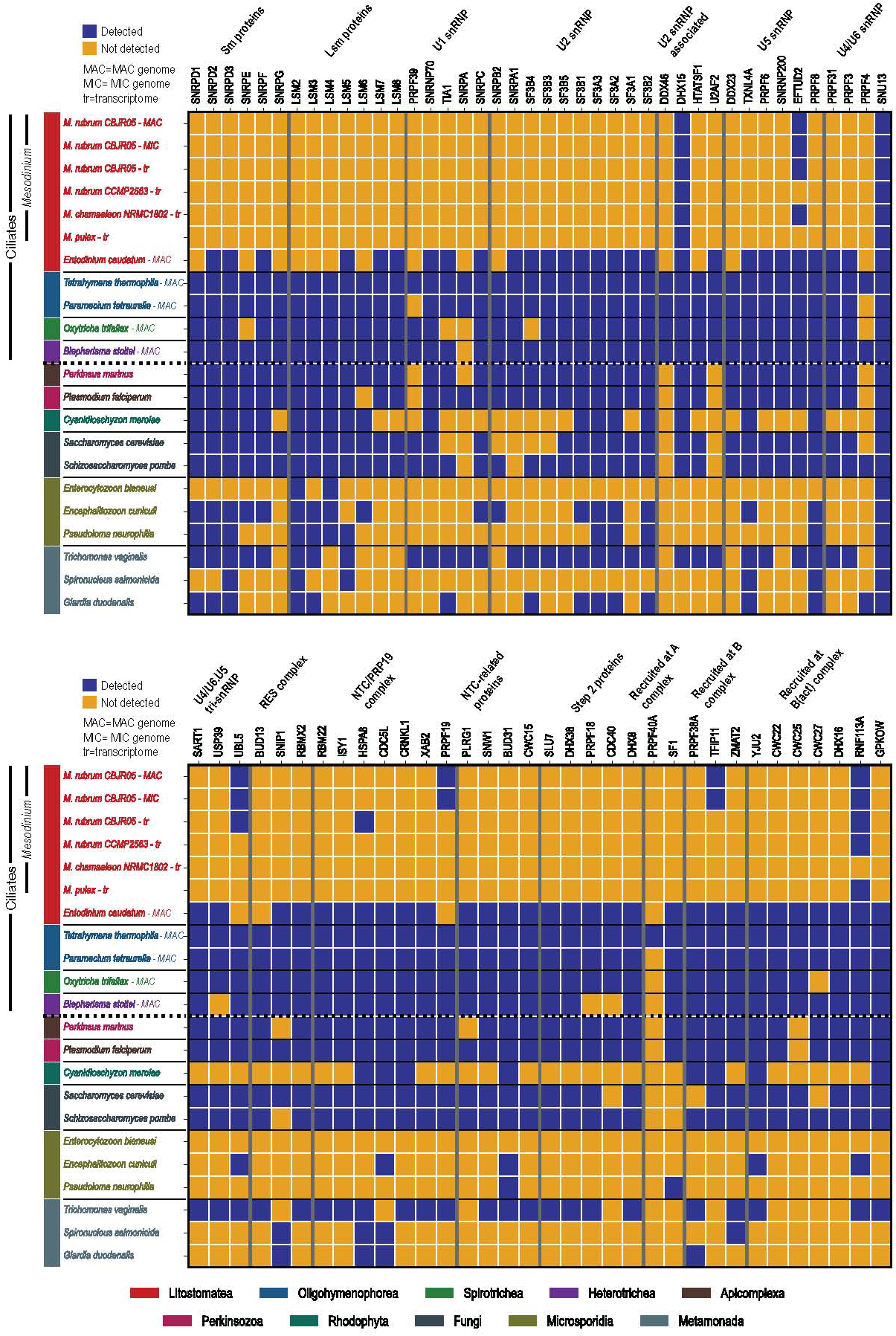
*Matrix of spliceosomal proteins detected in* Mesodinium rubrum*, other* Mesodinium *species, and relevant protists by OrthoFinder*.

Most striking among the spliceosomal proteins was the complete absence of all the RNA-binding, torus-forming Sm and Lsm proteins^67^ in *M. rubrum* (6 and 7 proteins respectively; BLASTP E-value < 1e-05). This contrasted with *T. thermophila*, in which all the Sm and Lsm proteins were detected with E-values ≤ 4e-15. The ciliate *Stentor coeruleus*, like *T. thermophila*, has an essentially complete set of spliceosomal protein orthologs^68^, despite possessing the shortest known spliceosomal introns^69^. BLAST hits to the *M. rubrum* proteome for the human-protein query sequences were unlikely to be orthologs as the search targets were not reciprocal (Data S1E).

Of the three putative spliceosome protein orthologs identified in *M. rubrum* by BLAST best reciprocal hits, two that were also consistently detected by OrthoFinder in all the *Mesodinium* species analyzed (DHX15 - gene g58807 and SNU13 - g40165) have additional, non-spliceosomal roles reported in the literature. DHX15 has been implicated in immunity against RNA viruses in association with the inflammosome protein NLRP6^70^. A DHX15 ortholog was also reported in the microsporidian parasite *Amphiamblys* sp., which has lost its introns and most spliceosomal proteins^19^. SNU13 has a well-known non-spliceosomal role in eukaryotes, interacting with C/D box snoRNAs such as the non-spliceosomal U3 snoRNA, serving in 2’O methylation and development of ribosomes^71^. *M. rubrum* possesses orthologs of other core complex proteins that associate with C/D box snoRNAs (i.e., Fibrillarin - g33303, NOP58 - g4333, and NOP56 - g9895). Furthermore, SNU13 is also present in the genomes of two intron-lacking microsporidian species (*Edhazardia aedis* and *Amphiamblys* sp.), both of which lack most spliceosomal proteins^19^.

The remaining putative spliceosomal protein ortholog in *M. rubrum* identified by BLASTP, but not OrthoFinder, has a weak match to the human U2AF2 query sequence (E-value 6e-8) but the reciprocal best hit of the human sequence is the ciliate one. In the *M. pulex* and *M. chamaeleon* transcriptomes, putative orthologs of both U2AF1 and U2AF2 were detected by reciprocal BLAST searches, but, like the putative ortholog in *M. rubrum*, their matches have much higher E-values than orthologs detected in *Tetrahymena* by BLAST. This suggests that the reciprocal best hits approach may have yielded false positive orthologs or that the proteins involved have diverged substantially from their conventional role.

### Absence of NMD proteins in Mesodinium rubrum

Given that we were unable to detect spliceosomal introns, spliceosomal noncoding RNAs (ncRNAs), and most spliceosomal proteins, we next searched for proteins involved in nonsense-mediated decay (NMD), which is proposed to detect introns that have erroneously failed to be spliced out; the pathway is generally conserved in eukaryotes including ciliates^72^.

The definitive NMD protein, Upf1, enables recognition of premature stop codons and degradation of mRNAs containing them^73^. The Upf1 homolog in *Tetrahymena thermophila* (TTHERM_00726300; UniProt Q24GG1) has characteristic N-terminal RNA helicase domains (Pfam: PF09416 and PF18141; amino acids 54-351; the remainder of the protein has non-specific AAA domains common in other proteins). We did not detect homologs of the *T. thermophila* Upf1 N-terminal domain region (E-value < 1e-5) in either *M. rubrum* nuclear genome assemblies using TBLASTN (search option:-db_gencode 29). This contrasts with heterotrich ciliates like *Stentor* and *Blepharisma*, which have tiny introns (predominantly 15 or 16 nt) and low intron densities^69,74^ (Table 1), but possess clear Upf1 homologs (e.g., one MAC-encoded *Blepharisma stoltei* Upf1 homolog has an E-value of 5e-67 vs. *T. thermophila* Upf1). We also did not detect homologs of *T. thermophila* Upf2 (TTHERM_00442760) and Upf3 (TTHERM_00046240) at the same E-value threshold. It is thus likely that the entire NMD complex has also been lost in *M. rubrum*, consistent with the role of NMD proteins in intron processing.

### Presence of rRNA ITSs, nonspliceosomal tRNA introns in tRNA-Tyr(GTA), and a eukaryotic tRNA splicing endonuclease

Previously, rRNA internal transcribed spacer (ITS) sequences were amplified in *M. rubrum* and other *Mesodinium* species, but, whereas it was proposed that the 5.8S might be fused to the preceding 18S region^28,75^, by means of Infernal searches^59^, we observed distinct 18S, 5.8S, and 28S subunits separated by ITSs in the *M. rubrum* MAC genome assembly (Data S1G). Besides ITSs, we observed the presence of additional ncRNA internal sequences that are spliced out during tRNA maturation: non-spliceosomal tRNA introns.

Thirteen tRNA^Tyr^ genes containing 13-nt non-spliceosomal introns were predicted in the *M. rubrum* MAC genome by tRNAscanSE; all had GTA anticodons, a high tRNA prediction score (74.7), and were identical except for their introns. The introns were nonetheless highly similar, with substitutions at only two nucleotide sites (IUPAC representation: AG[H]G[W]TCGCAGAA). No other tRNA introns were observed, particularly in putative tyrosine tRNA genes that permit translation of the reassigned *M. rubrum* stop codons, UAA and UAG.

In a mechanism distinct from spliceosomal intron splicing, tRNA introns in other organisms are excised by tRNA splicing endonucleases (TSENs)^76^. The *M. rubrum* MAC genome encodes five TSEN genes (Supplementary Results SR2). tRNA introns seemingly could be lost by deletion more easily than spliceosomal introns, since they only depend on one or two TSENs instead of the numerous proteins and ncRNAs of the spliceosome. However, this is not the case due to the role of these tRNA introns in the maturation of tyrosine tRNAs. The nature of these tRNAs, their modification, introns, and relation to the evolution of the unique *Mesodinium* genetic code in which two stop codons have been reassigned to tyrosine are further discussed in Supplementary Discussion SD3.

### Internal eliminated sequences (IESs) are enriched in the MIC genome assembly

Most ciliate MIC genomes are characterized by the presence of IESs, which are excised during MAC genome development^47,55,77–79^. Many, if not most, of these are transposon-derived; consequently, most commonly they are bounded by TA-repeats characteristic of the transposons from which the domesticated transposases arose^47,55,77–79^.

A total of 21,906 and 6,852 IESs were respectively predicted from the MIC and MAC PacBio HiFi reads mapped onto the MAC reference assembly, with mean retention scores of 0.54 and 0.21 (Figure 4; Figure S5). The latter suggests the MAC library contains about 21% MIC contamination, although most reads were of MAC origin. Retention scores of IESs predicted in the MIC library had a sharp peak at 1.0. Scores were lower on average for longer IESs (Figure S5B), indicating that the IES retention score in the MIC library was underestimated due to longer IESs being incompletely covered by reads. For subsequent analyses, we used a subset of 12,228 IESs predicted from MIC reads with retention score > 0.5, of which 7,591 (62%) were bound by tandem direct repeats containing a TA-submotif.

**Figure 4.**
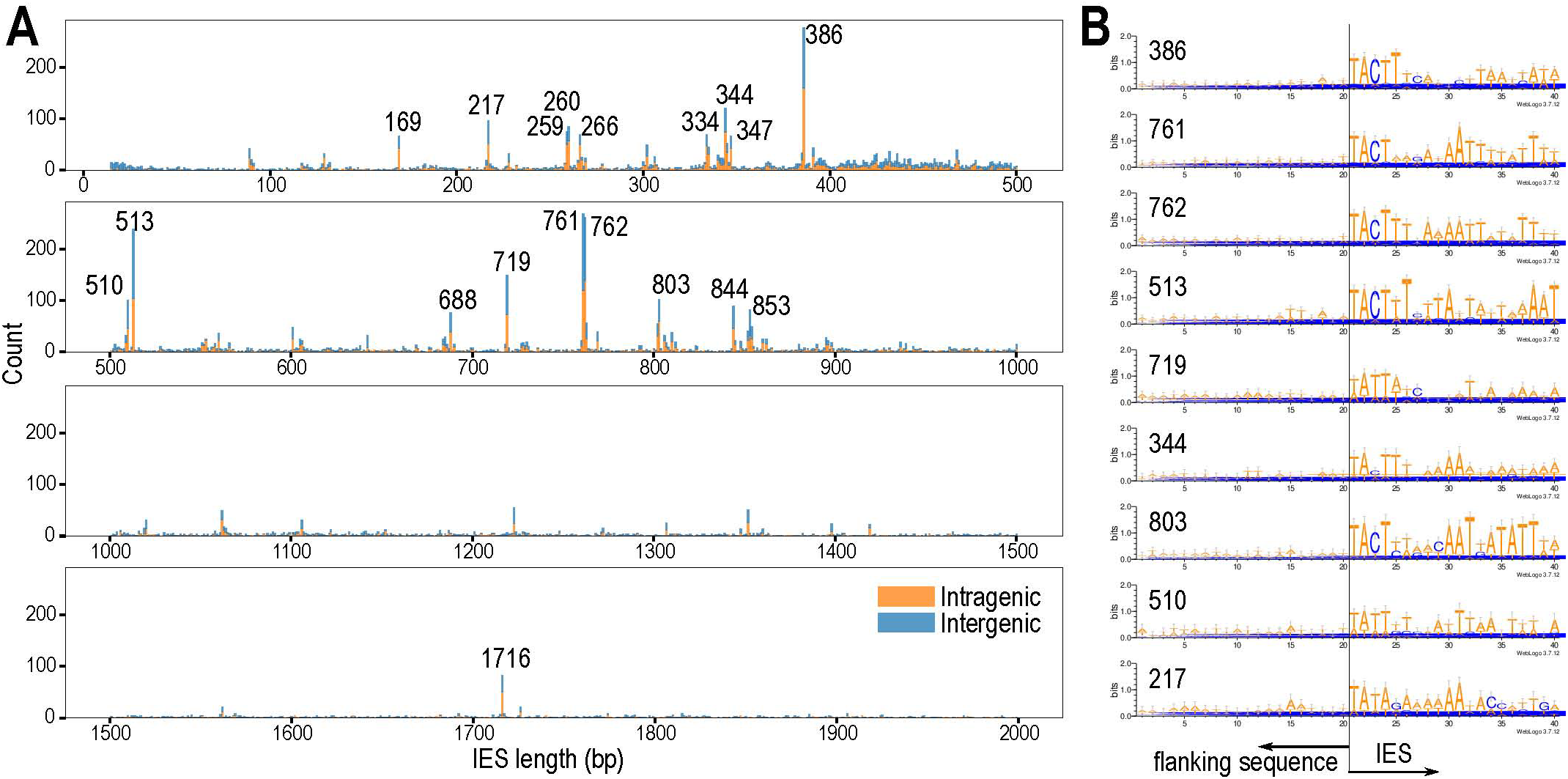
*Length distribution of* Mesodinium rubrum *IESs.* (**A**) Length histogram for IESs predicted from mapping MIC reads to MAC assembly, where retention score >0.5; peaks of abundant size classes are labelled. (**B**) Sequence logos at IES boundaries for size classes represented by ≥100 IESs, sorted by abundance.

Most *Mesodinium* IESs were several hundred bp long (mean 827 bp, median 707 bp). Like *Tetrahymena*^77^, but unlike *Paramecium* or *Blepharisma*^78,79^, there were few short (< 100 bp) IESs and there was no periodic length distribution. The IES length distribution (Figure 4) showed pronounced peaks at specific length classes; within each class, the sequence boundaries had conserved motifs when aligned with respect to the TA-submotif, consistent with them representing distinct families of repeat elements (top five by abundance: 386, 761, 762, 513, and 719 bp). About 51% of IESs (6,235 of 12,228) were located within predicted protein-coding genes, somewhat more than expected if IESs were uniformly distributed in the genome, given the overall coding density of 41% (77.6 Mbp), but this may be because low-complexity repeat regions were masked before IES prediction.

### Few potential horizontal gene transfers to *M. rubrum* despite karyoklepty

Using searches against a phylogenetically diverse reference database supplemented by ciliate and cryptophyte genomes, we found no evidence for extensive HGT between cryptophytes, particularly *T. amphioxeia*, and *M. rubrum*. Nevertheless, putative HGT candidates were identified, including a putative PAP/Fibrillin protein of apparent cyanobacterial or stramenopile origin and two copies of a cryptophyte-derived protein family of unknown function (Data S1H). The candidate HGT gene of most interest is of alphaproteobacterial origin and encodes an Npt1/Npt2-family nucleotide transporter (gene g4460). This gene was present in both MAC and MIC genome assemblies and the transcriptome; it contains reassigned stop codons, translated as tyrosine in *Mesodinium*’s nonstandard nuclear code, and thus is unlikely to be an environmental contaminant. All other genes on the 37 kbp MAC contig (contig_10838, 83x coverage) containing this gene appear to use *Mesodinium*’s genetic code. Genes flanking g4460 do not have any hits to known proteins in NCBI’s NR database, excepting g4457 and g4458, which have various hits to eukaryotes (∼45% identity) and *Chloroflexota* bacteria (∼25% identity), respectively. Homologs were detected in transcriptomes of another *M. rubrum* strain and *M. chameleon*, but not of *M. pulex*. Given that homologs were present in two *Mesodinium* species and adopted their genetic code, this indicates a relatively old acquisition.

The *Mesodinium* sequences fall within an alphaproteobacterial clade of Tlc-related proteins (Figure 5). Nearly all other members of the tree are putative plastidic ATP/ADP transporters from eukaryotic algae. Such a gene could enable energy transfer between the captured algal chloroplasts and host cells (Supplementary Discussion SD2). Our phylogenies indicate that several duplications or independent gains have occurred across eukaryotic lineages, which have been previously studied^80^. The top BLAST hit in NCBI’s NR database with an identifiable species belongs to *Candidatus* Megaira venefica (accession numbers: WP_322777370.1 and NZ_JARJFB010000143.1). This sequence is also the sister to the *Mesodinium* homologs in our phylogenies. *Ca.* Megaira venefica belongs to the alphaproteobacterial lineage *Rickettsiales*^81^ and is an endosymbiont of *Paramecium nephridiatum*, a brackish water ciliate^82^. Members of *Holosporales* and *Rickettsiales* are well-known endosymbionts of various ciliates^83–87^. The best BLAST hits in NCBI’s NR database within the genus *Rickettsia* are annotated as ATP/ADP exchange transporter Tlc1.

**Figure 5.**
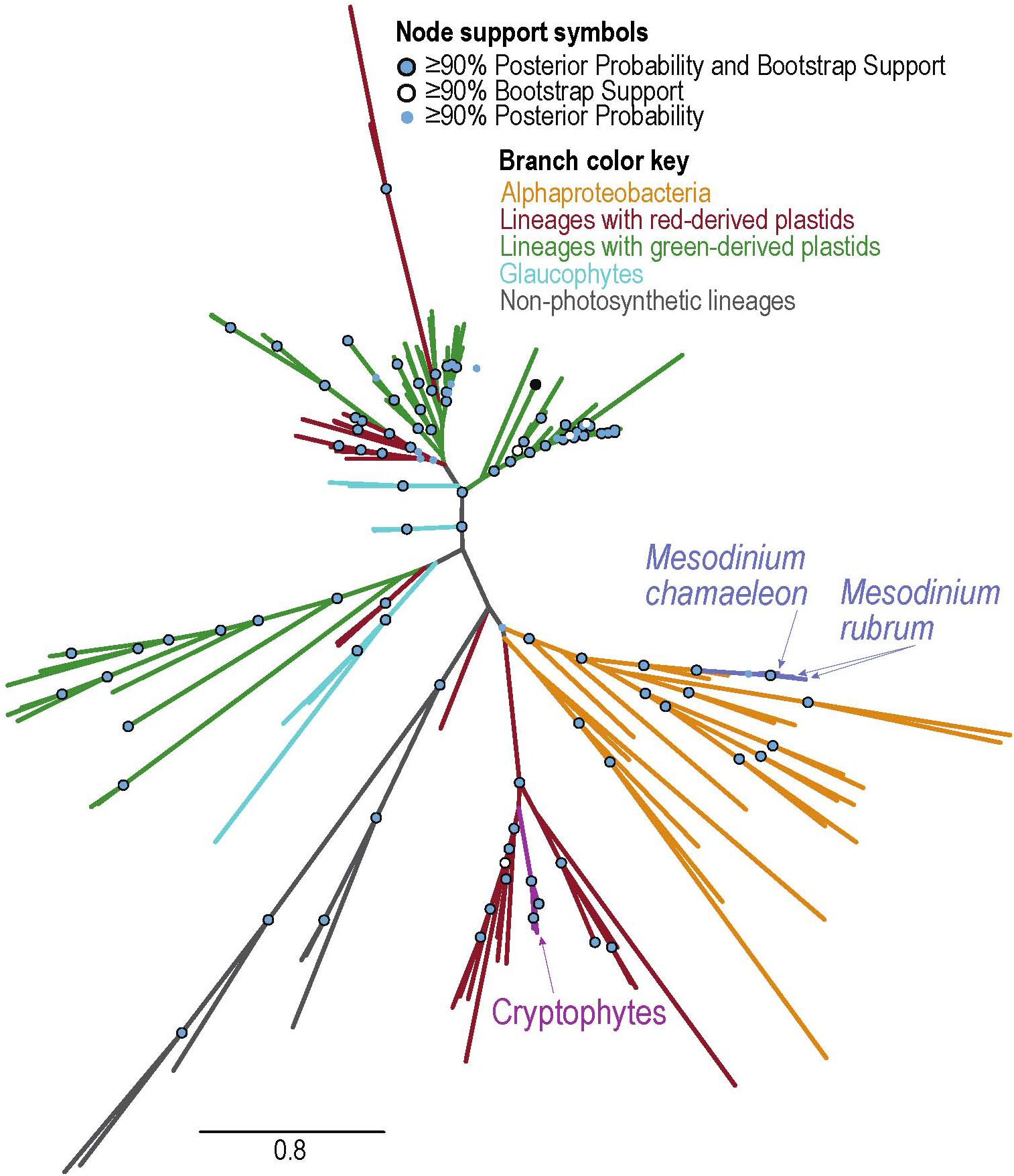
**Maximum a posteriori phylogeny for putative ATP/ADP transporter inferred under the LG+I+G4 model.**

## Discussion

The ciliate genus *Mesodinium* has long been problematic for taxonomy, due to its members’ reduced or divergent morphology^88^ and highly divergent rRNA genes^29^. Members of the *M. rubrum* species complex practice a highly specialized form of kleptoplasty, accompanied by theft of one or more transcriptionally active nuclei (karyoklepty) and almost total reliance on the metabolism of their prey for their energy needs^34^. Our analyses of the *M. rubrum* genome reveal further distinctive molecular peculiarities, namely a paucity of genes gained by horizontal transfer and the apparent absence of spliceosomal introns.

Extensive gene losses appear to have shaped *Mesodinium* genome evolution greatly. Previously, we reported substantial metabolic gene losses in the genus *Mesodinium* based on transcriptomic analyses, most pronounced in *M. rubrum*, including a striking absence of most peroxisomal genes^39^. Our current analyses of the *M. rubrum* nuclear genomes now indicate that introns are also absent, alongside virtually complete loss of the spliceosomal complex comprising over a hundred distinct ncRNAs and proteins, and the associated mRNA nonsense-mediated decay (NMD) pathway. Unlike the metabolic gene losses previously reported, which correlate with the shift from heterotrophy to near-complete metabolic dependence on captured organelles, the spliceosome losses appear to have occurred early in *Mesodinium* evolution.

Rather than thinking of intron loss as a return to a “primitive” state, *Mesodinium* should be seen as having re-evolved a robust system with less bureaucratic overhead. The loss of the RNA-splicing system is juxtaposed with the persistence of DNA splicing in the form of IESs. We think there are important reasons for this, outlined below. Furthermore, the costs of RNA splicing are likely far greater, given that they are continuously levied, whereas DNA splicing costs are levied only intermittently, during comparatively infrequent ciliate sex.

Not having spliceosomal introns can be advantageous to eukaryotes: (i) they become impervious to Introner mobile elements, which no longer have sanctuaries to lurk in, and thereby avoid intron blooming and the associated costs; (ii) they become impervious to intron-possessing mobile elements and viruses such as mirusviruses^89^; (iii) their transcription and translation become streamlined; (iv) they avoid the additional energetic cost of transcribing additional RNA, problems caused by splicing errors, and have a diminished need for NMD’s error-handling capabilities (on such costs, see reference 97). The first two advantages may also be shared by eukaryotes that can only efficiently splice short introns, e.g., most ciliate species and nucleomorphs. Though Introners were found to be most common in aquatic lineages, they were not reported in the ciliate species analyzed^9^. Eukaryotes with few introns, including some parasites, will also benefit from more efficient translation and a much-diminished need for NMD. Despite this, only *Mesodinium* and a few eukaryotes with highly streamlined genomes completely lack spliceosomes and spliceosomal introns.

### Hurdles posed by introns to horizontal gene transfers from algae in host ciliates

Among organelle-sequestering organisms, *M. rubrum* has a singular ability to regulate and replicate organelles stolen from *Teleaulax* or *Geminigera* cryptophyte prey, and to exploit prey metabolism, through karyoklepty (nucleus stealing)^31,34,90^. The stolen nucleus, which does not divide in *M. rubrum*, remains transcriptionally active and docks with the ciliate’s nuclei, apparently using the ciliate’s protein targeting machinery to control the various sequestered organelle complexes (SOCs)^31–34^. Because of how intimately the prey nucleus is exploited, it hypothetically has the potential for horizontal gene transfer (HGT). However, we observed no evidence for substantial HGT from *T. amphioxeia* or other cryptophyte algae to *M. rubrum*. A ratchet-like mechanism for organelle evolution^43^, in which horizontal gene transfers to the ciliate nuclear genomes are retained whilst their source genes are lost from the algal genomes, is precluded by *M. rubrum*’s regular replenishment of nuclei from *T. amphioxeia*^31^. On the other hand, despite the stable transgenerational transmission of their eukaryotic algal symbionts, algae-to-host HGTs have not been reported yet in the ciliates *Paramecium bursaria*^91^ and *Stentor pyriformis*^92^, though bacterial HGT to the former has been reported^93^.

Algal-to-host HGT may be rare simply because the ciliates cannot splice out algal introns effectively. *M. rubrum*, lacking a spliceosome, is faced with a mean intron density of 4.2 per gene in the genome of its prey, *T. amphioxeia* (Table 1). Other ciliate species that host algae have spliceosomes, but their own introns are typically very short (< 25 nt in *P. bursaria*^91^, mostly 15 nt in *S. pyriformis*^69,74,92^), with narrow length distributions that suggest their spliceosomes are structurally constrained and would not effectively splice the longer algal introns (and demonstrated for *P. bursaria*^94^), with the likelihood of splicing failure compounding for algal genes with multiple introns. A related argument has been made as to why nucleomorphs have not yet been lost: presuming that their tiny introns cannot be spliced out by host spliceosomes, transfers of intron-containing genes to the host cell nucleus would be non-functional^95^. Whatever the main mechanisms that hinder HGT may be (additional possible hindrances given in Supplementary Discussion SD7), they have not been completely prevented in *M. rubrum*’s lineage, as exemplified by the putative ATP/ADP transporter HGT we detected.

Fundamental differences in gene organization between eukaryotes and bacteria are likely a major impediment to successful bacterial gene transfers to eukaryotes: amongst others, persistence requires chance possession of suitable eukaryotic promoters for transcription, and also the absence of cryptic, coincidental introns that would delete essential genic regions. For inter-eukaryotic HGT, more consideration needs to be given to the role of possible incompatibilities in gene organization, like those between the intronless *M. rubrum* and intron-rich *T. amphioxeia*.

### Gains and retention of spliceosomal introns

Numerous *ad hoc* proposals exist for the origin of new introns, which may derive from internal duplications, inexact double-strand break repair, organellar DNA, or even exonic sequences; what these mechanisms have in common is that they generate sequences that fortuitously contain splice sites, and are recognized by the spliceosome as intronic^5–7^. More recently, it has been shown that mobile elements are not just sporadic sources of introns but also that certain transposons, collectively referred to as Introners, are responsible for *en masse* intron proliferation events across diverse eukaryotic lineages^8,9^.

Introns may persist in genomes through a neutral, ratchet-like mechanism: introns do not disrupt gene function because the spliceosome removes them from transcripts, but a mutated sequence that cannot be spliced is likely deleterious^96^. Thus, while intron gain is fairly easy, loss is difficult. Ironically, an efficient spliceosome hence encourages the persistence and accumulation of introns and predisposes a genome to keep acquiring them. In turn, having more introns necessitates the upkeep of an efficient spliceosome. Alternatively, intron acquisition and maintenance can be interpreted through a population-genetic lens, in which gains and retention of deleterious introns result from small effective population sizes^97,98^.

In general, introns are expected to be energetically costly, to enhance the production of problematic transcripts associated with splicing errors, and also increase the vulnerability of their allelic carriers to deactivating mutations (associated with splice-site motifs). Thus, in populations with very large effective sizes, selection will encourage loss-of-intron alleles. With introns also arising by various mechanisms, a stochastic steady-state equilibrium number of introns per gene (intron density) is expected. The equilibrium intron density will be reduced in populations with large effective sizes, where selection is more efficient. Unless this equilibrium density is very tiny, the spliceosome has a built-in guarantee for permanent existence, as it will be indispensable until the final intron in a key gene is physically deleted.

Ciliates possess another class of mobile element derivatives unique to them that they excise during development, but from germline DNA rather than pre-mRNA transcripts: IESs. Contrary to the popular characterization of IES excision as an adaptive defense mechanism, we interpret their evolution and persistence through a similar neutral, ratchet argument, with IESs, unlike introns, largely not being subject to selection because they are restricted to the germline^78,99,100^. In *M. rubrum*, processes that give rise to new IESs may have been active recently and may have contributed to genome size expansion after spliceosome loss (Supplementary Discussion SD8).

### The bases for large-scale spliceosomal intron loss

Given the logic of the evolutionary ratchet with its built-in irreversibility, how can introns, much less the spliceosome itself, ever be lost? Below, we consider explanations that have been offered for large-scale spliceosomal intron losses, as well as our own observations about tiny introns in ciliates and other organisms, and how they relate to *Mesodinium*. Multiple, potentially superimposed explanations may be valid for intron loss events in different eukaryotic lineages.

### Molecular mechanisms for intron loss

The simplest explanation for intron loss is selection against introns, which requires alternate, intronless or splicing-incapable alleles to be present in the population or to arise by mutation^98^. Introns may be rendered inactive or exonized through point mutations of intron splice sites^101^, with their remnants tolerated as long as they do not cause premature stops or disrupt critical functional domains. In principle, particularly for short introns, exonization by splice site substitutions may be much more rapid than intron deletion, since rates of single base substitutions are much higher than deletions of the size necessary to remove introns^102,103^.

An alternative hypothesized molecular mechanism to point mutations is reverse transcriptase-mediated intron loss (RTMIL), where reverse-transcribed mature mRNA homologously recombines with its corresponding genomic locus, resulting in the loss of introns, particularly from the 3’ end of the gene^104–107^. Most (but not all) ciliate genomes exhibit 5’-locational bias of introns in genes, a predicted signature of RTMIL (Figure S4), and retrotransposons encoding reverse transcriptases are abundant in some ciliate genomes including *M. rubrum* (Table S2, Table S3).

Retrotransposons, however, are extremely common in eukaryotes, so their abundance cannot be the sole reason for large-scale intron losses, while 5’-located introns may be preserved because they have functional roles, such as intron-mediated enhancement of gene expression^108^. Furthermore, the ciliate germline-soma duality poses a barrier to RTMIL, since most reverse transcription will not occur in transcriptionally inactive MICs but in MACs and their transcriptionally active developing precursors. Consequently, the reverse transcribed DNA products along with their RTases (which also possess integrase domains) will need to find their way out of MACs and into MICs in order to integrate and stably persist across sexual generations.

### Population-genetic and other factors that may influence intron loss

Most eukaryotes with extremely few or zero introns are parasites or pathogens with reduced genome sizes. However, this may reflect sampling bias, as eukaryotic genome sequencing has been historically biased towards those with small genomes for practical reasons. We do not think there is a necessary causal link between trophic lifestyle or genome reduction and intron loss, due to the numerous counterexamples (Table 1, Supplementary Discussion SD9); many parasites and pathogens have large genomes and/or numerous introns. Genome size is not an especially reliable predictor of intron density either. Intron densities of the two yeasts in Table 1 differ by more than an order of magnitude, despite similar-sized genomes. More striking, the 0.37 Mbp nucleomorph genome of *Bigelowiella natans* has 852 predicted introns^27^, while the 180 Mbp genome of *Trichomonas vaginalis* has just 63^109^.

Nonetheless, selective factors that favor genome reduction may also favor the streamlining and loss of spliceosomal machinery (Supplementary Discussion SD10), as seen in the multiple losses of splicing in microsporidia, parasites with reduced genomes^110^. Furthermore, genomes are evolutionarily dynamic, and the sequences we observe are a snapshot of evolutionary history. Large genomes with few introns may be descended from ancestral genomes that underwent reduction and concomitant large-scale intron loss, followed by genome expansion without intron gain.

Population-genetic models of intron evolution via mutations that affect splice sites propose that large effective population sizes favor intron loss by purifying selection, whereas intron gains are more likely to drift to fixation in populations with small effective sizes, despite weak selective disadvantages^97,98^ (Supplementary Discussion SD11). Although population-genetic parameters have not previously been estimated for *Mesodinium*, *M. rubrum* is globally distributed^28^ and can form immense blooms spanning hundreds of kilometers, so very large effective populations are not implausible (Figure 1B; Supplementary Results SR6 estimates the *M. rubrum* effective population size given mutation rates of other ciliates). Moreover, *M. rubrum’*s cryptophyte prey must have comparable peak abundance, yet they have plenty of introns and no apparent significant intron loss (Table 1; Supplementary Discussion SD11). In the future, the population genetics of *Mesodinium*, *Teleaulax*, and other cryptophytes, including their mutation rates, needs to be carefully investigated to ascertain how much this may influence intron densities.

Introns in ciliates are notably short compared with other eukaryotes: the shortest known introns (15 nt mode) belong to heterotrich ciliates^69,74^, with their sister clade, the karyorelicts, not far behind (17 nt mode)^49^. This can influence intron loss/gain in two ways: (1) tight, short intron-length distributions may indicate a structurally constrained spliceosome that is incapable of splicing longer introns, which may not only be a hurdle to HGT (see above) and to the origin of new introns (since the available sequence space is smaller), but also leave little room for introns to encode secondary functions, e.g., regulatory elements that interact with promoters; and (2) short, 3n introns would be less likely to cause premature stop codons if erroneously retained, and would result in insertions of only a few amino acid residues; the probability of premature stops is also reduced by alternative genetic codes with stop-codon reassignment (Supplementary Discussion SD12).

### Possible scenarios for intron losses in Mesodinium

#### Scenario 1 – Gradual loss by intron-to-exon conversion and prevention of new intron acquisition

As described above, a conventional population-genetic explanation may partly account for intron losses in the genus *Mesodinium*. If the rate of intron loss outweighed intron gain, introns could have been gradually lost until none were left (Figure 6). Plausible molecular mechanisms in the *Mesodinium* ancestor include reverse-transcriptase-mediated intron loss (RTMIL) and exonization of short introns.

**Figure 6.**
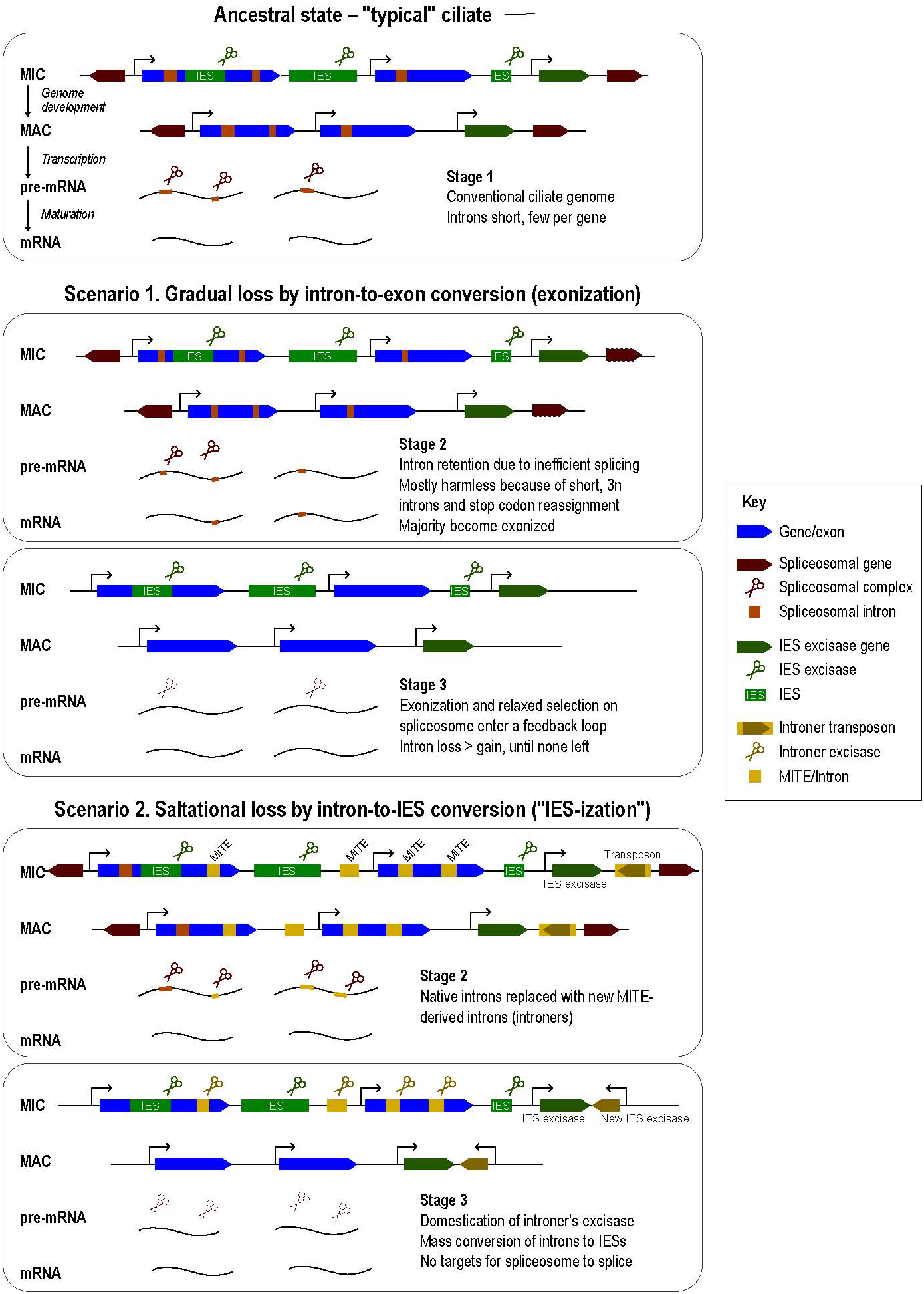
**Gradual vs. saltational scenarios for loss of spliceosome and its introns in the Mesodinium *ancestor*.**

We propose that exonization was favored in a *Mesodinium* ancestor having few, short introns per gene, with several factors lowering the cost of erroneous intron retention. Importantly, if its spliceosome had a narrow preference for short, 3n introns, like heterotrichs today, combined with its genetic code having only one stop codon, most intron retentions would not lead to frameshifting or premature stops but insertions of only a few amino acid residues. If some mutation in a spliceosomal gene now caused a slight decrease in spliceosome efficiency, this would select against deleterious introns (either longer, stop-containing or frameshifting), hastening exonization or promoting intron-free alleles. By further relaxing purifying selection on spliceosome efficiency, this would start a feedback loop.

The argument against this “gradual loss” scenario is that other ciliates do have short introns and 5’-locational bias but have not lost their introns completely. Loss may result in low intron density, but those that remain become asymptotically harder to get rid of, according to the ratchet-like evolutionary processes described above, as exemplified by other eukaryotes with extreme intron loss, like *Cyanidioschyzon*, which still maintain a functional, albeit reduced spliceosome to deal with only a handful of intron-bearing genes. Speculatively, the *Mesodinium* ancestor may have experienced pronounced genome reduction, as suggested by general gene losses^39^, and concomitantly reached a tipping point of intron numbers below which loss was more strongly favored, but later expanded its genome again without re-acquiring spliceosomal introns (Supplementary Discussion SD10).

#### Scenario 2 – Saltational loss by intron-to-IES conversion

Ciliates already have a mechanism that removes junk DNA, namely IES excision. Could introns be lost by conversion to IESs? We think that such conversion is unlikely for gradual loss of introns. The majority of *M. rubrum* IESs (Figure 4) are longer than the typical litostome ciliate introns (20-75 bp, Figure 2) so do not appear to represent converted introns. The most common boundary motifs of ciliate IESs (TA-TA) and introns (GT-AG) do not coincide. Thus, introns converted to IESs would likely not be excised at exactly the same boundaries, and would need to meet several criteria simultaneously, i.e., avoidance of excessive deletion of essential coding sequences, addition or removal of few, nonessential codons, and preservation of the translation frame; this would be improbable.

Instead of a gradual loss, intron-to-IES conversion could have occurred in a single episode, possibly within a single sexual generation (Figure 6). First, the *Mesodinium* ancestor with conventional spliceosomal introns is invaded by a novel Introner element that rapidly replaces all or most of its original introns in a blooming event, like in the chordate *Fritillaria*^111^. Soon afterwards, the transposase associated with this Introner is domesticated as a genome-editing excisase by coming under the control of a host promoter. The new introns are thereafter recognized as IESs and excised by the genome editing machinery, and no longer present in the somatic MAC genome; given that gene expression in ciliates is largely limited to the MAC, they are hence also absent from mRNAs. For excision to happen cleanly without any footprint, the Introner should be of a type with precise excision, and where either one of the splice sites or both are co-opted from the tandem sequence duplications (e.g., types B, C, or E in Figure 2 of Gozashti et al., 2022). The spliceosome, having lost its raison d’etre, is lost, making this intronless state permanent. Over time, genetic drift of the Introner/IESs and new waves of mobile-element invasion obscure any traces of what had happened. Because the Introner-IESs originate from a single transposon family, they are indistinguishable from other transposon-derived IESs in the genome.

Rapid replacement of old introns by an Introner element is key to this scenario because the splice sites of these new Introner/IESs must be sufficiently intact to be recognized by the domesticated transposase, whereas these motifs are eroded by genetic drift in older introns. In *Fritillaria*, the noncanonical introns are diverse, but individual families have highly conserved terminal repeats. Perhaps *Mesodinium* was invaded by only a single Introner family. Additionally, these Introners may have attracted suppressive epigenetic marks, which in ciliates can also help recruit the IES excision machinery, further encouraging their excision. Domestication of new excisases by a ciliate is not implausible; although PiggyBac family excisases are most frequently the primary ones for genome editing, there are exceptions in spirotrich ciliates like *Oxytricha*, which must have been independently acquired after the common origin of genome editing in ciliates^55,74,112^.

## Materials and Methods

Unless otherwise indicated, chemical reagents were analytical grade and purchased from Sigma-Aldrich or Merck, and procedures performed at room temperature (20 to 24 °C).

### Cultivation of Mesodinium rubrum CBJR05

*Mesodinium rubrum* strain CBJR05 was originally isolated from the James River in Chesapeake Bay in 2011^113^. *Teleaulax amphioxeia* GCEP01 was originally isolated from Eel Pond, Falmouth, Massachusetts in 2008 by Mengmeng Tong^114^.

*Mesodinium* was cultivated in 35 g/L artificial seawater prepared from commercial mix (hw Marinemix reefer, Wiegandt GmbH), supplemented with F/2-Si trace elements (CCAP protocol: https://www.ccap.ac.uk/wp-content/uploads/MR_f2Si.pdf)^115^. *Teleaulax* was cultivated in natural seawater from Helgoland, 32% salinity, supplemented with F/2-Si. Media were sterilized by filtration (0.2 µm). Both cultures were incubated at 18 °C on a 14 h light / 10 h dark cycle at 24 µmol photons m^-2^ s^-1^. *Mesodinium* stock cultures were subcultured (1:5 dilution into fresh medium) every ca. 10 days, and supplemented with dense *Teleaulax* culture (0.5 mL per 100 mL *Mesodinium*) when judged necessary based on inspection of algal density. *Teleaulax* was subcultured every 10 days (1:40 dilution into fresh medium). For larger cultures, volumes were doubled once per week, adding *Teleaulax* as needed.

### Immunofluorescence of M. rubrum for microscopy

Dense *M. rubrum* CBJR05 culture was centrifuged in pear-shaped flasks with an oil-testing centrifuge (280 g; 2 min). Concentrated cells were gently transferred by plastic Pasteur pipettes to microcentrifuge tubes in 1.5 mL aliquots. To each aliquot, 150 µL of 20% w/v formaldehyde stock solution was added to fix cells (>5 min), which were then centrifuged (1000 g; 1 min) and resuspended in 600 µL of bovine serum albumin (3% w/v) dissolved in TBSTEM buffer (10 mM EGTA, 2 mM MgCl_2_, 10 mM Tris-HCl, 1% v/v Tween-20, pH 7.4) (hereafter BSA/TBSTEM) to quench fixation. After 10 min, each aliquot of fixed cells was centrifuged and resuspended in 1 mL BSA/TBSTEM and stored at 4 °C until use.

Antibodies were diluted 1:100 or 1:200 v/v in BSA/TBSTEM (Table S6). Fixed cells were successively centrifuged (500 g; 1 min) and resuspended in: 100 µL of diluted primary antibody (30 to 60 min), 400 µL BSA/TBSTEM (wash, 5 to 15 min), 100 µL secondary antibody (30 min), 400 µL BSA/TBSTEM with 1 µg/mL DAPI (counterstain). A sample was inspected under epifluorescence to confirm antibody labeling. Finally, cells were resuspended in 60 µL ProLong Gold mounting medium, coverslipped (20 µL per slide), and cured overnight before imaging.

### Immunofluorescence and flow sorting of *M. rubrum* nuclei

Dense cultures of *M. rubrum* CBJR05 were concentrated by centrifugation as described above and gently transferred with a plastic Pasteur pipette to a 15 mL centrifuge tube. Concentrated cells were briefly centrifuged to remove waste debris (50 g; 1 min), supernatant with cells was transferred to a new tube, then centrifuged again (500 g; 2 min). The cell pellet was resuspended to 1:100 original culture volume in ice-cold Galbraith’s solution^116^ (45 mM MgCl_2_, 30 mM sodium citrate, 20 mM MOPS pH 7, 0.1% w/v Triton X-100, filtered at 0.22 µm pore size, stored at 4 °C until use), and pipetted up and down 10 times to lyse cells. Lysate was stained with DAPI (final conc. 1 µL/mL) for 5 min on ice; a sample was inspected under epifluorescence microscopy to confirm that nuclei were well separated. Lysate was fixed with formaldehyde (added 4% w/v stock to final conc. 0.2% w/v) overnight at 4 °C.

Fixed nuclei were centrifuged (1000 g; 2 min for all steps). To quench fixation, the pellet was resuspended by pipetting in 500 µL 3% BSA/TBSTEM and kept for 10 min. Nuclei were centrifuged and resuspended again in 250 µL BSA/TBSTEM to wash. Antibody labeling was performed as for whole cells except that reagent volumes were halved, and after DAPI staining, nuclei were resuspended in ice-cold Galbraith’s solution and held on ice until flow sorting.

The immunofluorescence-labeled nuclei suspension was sorted by fluorescence-activated cell sorting (FACS) on a BD FACSMelody instrument (nozzle 100 µm, pressure ca. 23 PSI, drop frequency 34.0 kHz, in “purity” sort mode), controlled with BD FACSChorus v1.1.18.0 software. Fluorescence settings for each dye were: DAPI – 405 nm excitation, 448/45 filter; Alexa Fluor 488 – 488 nm excitation, 507LP mirror, 527/32 filter. PMT voltages were: forward scatter 460 V, side scatter 490 V, DAPI 440 V, Alexa Fluor 488 590 V. Exact PMT voltages and gating parameters were interactively adjusted per run depending on the scatter plot. *M. rubrum* MAC and MIC nuclei, as well as *Teleaulax* kleptokarya, were gated according to the scheme in Figure 1E, and sorted into 1.5 mL microcentrifuge tubes (Eppendorf Lo-Bind). Sorted nuclei were centrifuged (5000 g; 2 min; 4 °C); supernatant was removed by pipetting; pellets were flash-frozen on liquid nitrogen and stored at-80 °C until use.

### Polarized light microscopy

*M. rubrum* or the dinoflagellate *Akashiwo sanguinea*, used as a positive control, were either sampled from dense cultures or concentrated using an 8 mm Transwell (Corning 3428) 6-well plate insert, and washed with F/2-Si media as described previously^113^. Cells were stained with 5-10× SYBR green (Thermo Fisher S7563) and mounted live in 1% low melt agarose on glass slides under coverslips. Cells were imaged on a Nikon Ti-E inverted microscope stand configured as an LC-PolScope, as described in (Mehta, 2013)^117^. Cells were viewed using a strain-free 60×, 1.2 NA water immersion plan apochromatic objective lens. Digital images were captured using a Hamamatsu Flash4 camera and processed using a Java-based image processing system for calculating images of differential retardation and slow axis orientations (OpenPolScope system^118^).

#### M. rubrum genome sequencing and assembly

DNA was extracted from sorted nuclei with the GenElute Mammalian Genomic DNA kit (Sigma Aldrich G1N70-1KT). Each batch (comprising between 200 k to 550 k nuclei) was digested with 180 µL lysis buffer T and 20 µL proteinase K (55 °C; 1 h), then diluted with 200 µL lysis buffer C. Formaldehyde cross-linking was reversed by heating at 80 °C for 1 h. Lysate was then column-purified following the manufacturer’s instructions and eluted in 100 µL elution buffer. DNA concentration was measured by fluorometry (Qubit DNA HS kit, Thermo Fisher Q33231).

For each nucleus type, 10 ng of genomic DNA (gDNA) was used for library preparation. MAC gDNA was not sheared, while MIC gDNA was sheared on a Meg3 at speed 52 (½ time). Libraries were prepared with the Pacific Biosciences CCS amplified ultra-low library kit (PacBio 101-980-000) and size-selected on a BluePippin (3.5 to 17 kbp for MAC, 4 to 17 kbp for MIC), and sequenced on a Sequel II instrument in CCS mode to yield 21 Gbp of HiFi reads per library.

The adaptor sequence “AAGCAGTGGTATCAACGCAGAGTACT” was trimmed from HiFi reads with bbduk.sh from bbmap v39.01^119^ with parameters: “ktrimtips=30 k=21 mink=15 overwrite=true”. Read libraries of the MAC and MIC sorted nuclei were assembled with Flye v2.9.2^53^ with options: “-g 300m --no-alt-contigs --scaffold --pacbio-hifi”. Co-assembled bacterial contaminants were identified by screening for rRNA genes and linked contigs. Sequences with >60% GC were excluded (10 Mb for the MAC and MIC genome assemblies), as the contaminants were only found in this GC range. No mitochondrial contigs were found, by alignment with a previously assembled mitochondrial genome of the same strain. Genome coverage estimates in Table S1 were obtained from the Flye output. The assembly version available from JGI’s Phycocosm web portal was further filtered to remove contigs <1 kbp with no annotated features; the original assembly is available at https://doi.org/10.17617/3.REWBDY.

Workflow: https://github.com/Swart-lab/mrub-sorted-asm

#### Targeted assembly of *M. rubrum* germline-specific sequence

To analyze sequences specific to or enriched in the MIC genome, PacBio HiFi reads from MIC and MAC libraries were separately mapped on the MAC genome assembly with minimap v2.24^120^ with parameter:-ax map-hifi. The mappings were converted to BAM format, sorted, and indexed with Samtools v1.15^121^, then analyzed with the MILRAA tool in BleTIES v0.1.11^122^ in “subreads” mode, (parameter: --type subreads), which identifies potential IES loci, clusters putative IES-containing reads, and performs a targeted local reassembly of the IES sequence, but which only can detect insertions, not deletions. Predicted IESs within low-complexity tandem repeats (below) were excluded, as the alignments may be less reliable.

Pipeline managed with Snakemake v6.8.1^123^: https://github.com/Swart-lab/mrub-sorted-compare

#### Annotation of repeats and low-complexity sequences in *M. rubrum* genomes

Interspersed repeat families in both M. rubrum de novo genome assemblies were predicted with RepeatModeler v2.0.4 (Table S2), then the repeats identified in the MIC genome were used to mask both the MIC and MAC assemblies with RepeatMasker. Low-complexity tandem repeats were annotated with TRF v4.09.1, using options “2 5 7 80 10 50 2000-d-h-ngs”; overlapping low-complexity tandem-repeat regions ≥1 kbp were merged with bedtools v2.30.0 (total 33,551,209 bp in MAC).

Workflow: https://github.com/Swart-lab/mrub-sorted-compare

#### M. rubrum transcriptome sequencing

400 mL of *M. rubrum* CBJR05 culture was starved until the density of free *Teleaulax* prey was judged to be minimal. Cells were concentrated by centrifugation (280 g; 2 min) in pear shaped flasks, gently transferred to 15 mL centrifuge tubes, then pelleted (500 g; 2 min), and supernatant was removed by pipetting. Each pellet was dissolved by adding 2 mL TRIzol reagent and pipetting up and down on ice, then split into 1 mL aliquots. To each aliquot, 200 µL chloroform was added, mixed by shaking for 2 min, then centrifuged (12,000 g; 15 min; 4 °C). Per aliquot, ca. 500 µL of the upper aqueous phase was transferred to new tubes, and mixed with 500 µL ethanol, then transferred to spin columns for purification with the RNA Clean & Concentrator 5 kit (Zymo Research) following manufacturer’s instructions. Each aliquot was eluted in 8 µL of elution buffer; RNA concentration and quality were checked by Nanodrop and gel electrophoresis. RNA-Seq libraries (two technical replicates) were prepared with the NEBNext Ultra II Directional RNA Library Prep Kit for Illumina (NEB E7760S), using the protocol for the NEBNext Poly(A) mRNA Magnetic Isolation Module (NEB E7490), and sequenced on an Illumina NextSeq 2000 instrument.

#### Screening of *M. rubrum* RNA-Seq data for potential introns

RNA-Seq libraries were mapped to the *M. rubrum* MAC and *T. amphioxeia* genomes with a modified version of HISAT2^124^, where minimum intron length was lowered to 9 (vs. default 20), to account for short introns in ciliates (https://github.com/Swart-lab/hisat2/commit/d88fbdd6). For consistency (Supplementary Discussion SD5), we mapped published RNA-Seq data from other litostome ciliates and eukaryotes with reduced spliceosomes (Data S1I)^15,17–19,57,58,125–130^ to available genome assemblies with the same mapper.

RNA-Seq mappings were processed with Portcullis v1.1.2^131^ to identify potential splice junctions and calculate metrics to evaluate whether these represent true introns (“prep” and “junc” steps). For ciliates, the “filter” step retained many short (about < 20 bp) junctions without canonical splice site motifs. Based on what is known about ciliate introns in model species, these were likely to be false positives, although their length range overlapped with true introns. We hypothesize that the metrics used by Portcullis to train its random forest model are less effective with ciliates because their true introns are short and have less available information.

We therefore compared splice junctions across different species empirically from the “junc” results, with the following metrics: length distribution, relative proportion of canonical vs. other splice site motifs, Hamming distance (a measure of repetitiveness in the neighborhood of the splice junction, more repetitive means more likely to be a mismapping).

Workflow: https://github.com/Swart-lab/mrub-intron-analysis

#### Screens for spliceosomal components

Searches for the U1, U2, U4, U5 and U6 spliceosomal RNAs used Infernal 1.1.5^59^ with default parameters and the Rfam 15.1 database^132^.

Table 1 of reference 63 was used as the basis for which spliceosomal proteins to search for in the *M. rubrum* MAC genome. We omitted a few yeast proteins without human orthologs, and replaced some accessions of human genes with their UniProt primary synonyms (Release 2025_03)^133^. As query sequences we used the proteome of *Tetrahymena thermophila* (version 6 from ciliates.org, file: Tetrahymena_thermophila_protein_v6.fasta), a ciliate with a complete, well-annotated proteome, and the human proteome from UniProt Release 2025_03.BLASTP and TBLASTN (version 2.9.0) searches for spliceosomal proteins used default parameters, except for the *Mesodinium* genetic code (table 29) (TBLASTN), and E-value threshold ≤1e-05 (following^19,68^). We searched both *T. thermophila* and *M. rubrum* for BLASTP reciprocal best hits with human sequences as the initial query.

Reciprocal best hit BLASTP searches for orthologs were also performed for de novo assembled transcriptomes from four *Mesodinium* species: *M. rubrum* CBJR05 (the reference strain used for genome assemblies), *M. rubrum* strain CCMP2563, *M. chamaeleon* strain NRMC1802, and *M. pulex* strain EPMP20B2, using proteins that were previously filtered for contaminants^39^.

For comparative analyses of spliceosomal protein components, we obtained proteomes from protists and other eukaryotes of interest and used OrthoFinder v3.1.0^134^ to predict orthogroups and orthologs (Data S1F). Our OrthoFinder analysis used the *Tetrahymena thermophila* SB210 strain from GenBank (accession GCA_000189635.1) with the reference list of spliceosomal components based on gene IDs from the *T. thermophila* macronuclear genome V6 annotation from the Tetrahymena Genome Database (https://tet.ciliate.org/), with mapping between these databases based on BLASTP best hits. Human spliceosomal proteins without orthologs detected in *Saccharomyces cerevisiae* were not included from Figure 3.

#### Gene prediction and annotation in M. rubrum

RNA-Seq was mapped to a combination of the *M. rubrum* MAC and to the JGI *T. amphioxeia* genome assemblies (JGI GOLD Project ID Gp0578332) with HISAT2 with the parameter “--min-intronlen 9”. Separate BAM files corresponding to reads mapping to either *M. rubrum* or *T. amphioxeia* were generated by extracting reads mapping to contigs of each.

After modifying AUGUSTUS source code to accommodate the *Mesodinium* genetic code and training on a manually curated gene set, gene prediction was performed using AUGUSTUS v3.5.0, parameterized for *M. rubrum*. Predictions were run in complete gene model mode with intron prediction disabled, incorporating RNA-seq–derived extrinsic evidence in the form of GFF3 hints generated from the RNA-seq BAM files. Soft masking was disabled, and gene models were constrained by a minimum intron length of 9 bp as specified during hint generation. Configuration files, modified source code and scripts are deposited at https://doi.org/10.17617/3.REWBDY.

Ribosomal RNAs (rRNAs) were predicted with Infernal v1.1.5^59,135^ using RFAM models for eukaryotic 18S, 28S, and 5.8S rRNA.

InterProScan (version InterProScan-5.75-106.0 with switch “--excl-applications ProSiteProfiles,ProSitePatterns,Hamap,PRINTS,MobiDBLite”) was run from the command line to annotate domains in the predicted proteins for the *M. rubrum* MAC and MIC genomes and *Tetrahymena thermophila* MAC genome gene predictions from ciliates.org (Tetrahymena_thermophila_protein_v6.fasta).

Pfam domain annotations from the InterProScan annotations of *M. rubrum* proteins from the MAC genome assembly were used to identify tRNA splicing endonucleases.

#### Distributions of synonymous substitution rates and 4-fold synonymous site diversity

Distributions of *M. rubrum* MAC coding sequence synonymous substitution rates were obtained with wgd v2.0.38^136^ source code was modified to accommodate the alternative genetic code of *M. rubrum*: (1) the universal (default) genetic code of PAML 4^137^ in “tools.c” was substituted with the *Mesodinium* genetic code before recompiling this software to create a fresh codeml binary (wgd v2 uses codeml to obtain K_s_ values^136^); (2) the parameter “table=29” was added to the BioPython “translate” function call in wgd v2’s core.py.

Sequence diversity estimates at 4-fold synonymous sites were calculated with a custom Python script that used the coding sequence alignments from wgd v2, restricted to genes within collinear genomic segments in the “segments_coordinates.tsv” output file produced by wgd v2.

#### Genome library preparation, assembly and gene prediction in *T. amphioxeia* GCEP01

Long-read sequencing libraries were constructed using a SMRTbell Template Prep Kit 3.0 (Pacific Biosciences), sheared using a Megaruptor (Diagenode) or a needle and sized on either a SAGE ELF or a Blue Pippin Instrument (Sage Science, Beverly, MA, USA). Sequencing was performed on a PacBio Sequel II in CCS mode using a 30 hr movie time with 2 h pre-extension.

A total of 69.90x PacBio CCS read coverage (mean read length 14,567) was assembled with HiFiAsm v0.16.1^138^ and polished using RACON v1.4.10^139^, yielding an initial 406.5 Mbp assembly (contig N50 61.1 kbp). The scaffolds were screened against bacterial proteins and organelle sequences using GenBank nr^140^. Homozygous SNPs and indels were corrected in the release by aligning Illumina fragment reads (150 bp paired-end, 400 bp insert, 33.0⨉ coverage) with bwa mem^141^ and identifying homozygous SNPs and indels with the GATK’s UnifiedGenotyper tool^142^. 2,038 homozygous SNPs and 18,069 homozygous indels were corrected in the consensus sequence. The final release consisted of 204.7 Mbp of sequence with a contig N50 of 123.4 kbp.

Raw RNA-Seq datasets for *T. amphioxeia* collected at mid-day or mid-night either under high (100 μmol quanta m^-2^ s^-1^) or low (10 μmol quanta m^-2^ s^-1^) light conditions were downloaded from the NCBI Sequence Read Archive (SRA) using SRA Toolkit. Raw reads were quality trimmed using Trim Galore version 0.6.10 with paired-end mode, NextSeq quality threshold of 25, and a minimum length of 60 bp. Post-trimming quality was assessed with FastQC, and trimmed reads were mapped to the JGI *T. amphioxeia* genome using STAR version 2.7.10b with minimum intron length set to 9 bp, with strand-specific intron information, and final file output as BAM sorted coordinate files^143^.

BRAKER3 3.0.8 with arguments “--augustus_args=”--min_intron_len=9“” was executed in a Singularity container (version 3.7.2+17) on the JGI *T. amphioxeia* genome using the STAR-generated RNA-seq BAM files and the JGI *T. amphioxeia* GeneCatalog proteins, clustered at 99% with CD-HIT^144^, as protein data^145,146^. Gene structure statistics were extracted with AGAT (agat_sp_statistics.pl) from the BRAKER3 gff3 file and the mRNA without isoform statistics were used^147^. The intron length mode was calculated using the GenomicRanges package^148^ in R version 4.5.1.

#### BUSCO analyses

BUSCO version 6.0.0^66^ with default parameters was used along with the alveolate lineage database “alveolata_odb12” to assess genome completeness (Table S4). The database used comprises 99 orthogroups and includes five ciliate species: *Tetrahymena thermophila*, *Blepharisma stoltei*, *Euplotes crassus*, *Halteria grandinella* and *Ichthyophthirius multifiliis*. Determination of which of the “missing” BUSCO database entries correspond to spliceosomal proteins was based on BLASTP searches of GenBank (nr_cluster_seq database) with individual sequences from alveolata_odb12 as queries.

#### Intron metrics in other ciliates

For ciliates where published gene predictions are available, we evaluated intron frequency and relative position within gene bodies, to see if these are consistent with RTMIL model of intron loss. Genome annotations were obtained from Ciliates.org and ParameciumDB, supplemented with the litostomes *Entodinium caudatum* and *Balantidium ctenopharyngodoni* (Data S1I).

Workflow: https://github.com/kbseah/ciliate-introns/

#### Horizontal gene transfer analyses

To look for potential HGTs, a database with representation across the tree of life was used^149^, supplemented with ten additional ciliate genomes, two cryptophyte genomes, and two transcriptomes (Data S1K). The location of each gene, its functional annotations, the distribution of its homologs throughout Ciliophora, its best-scoring BLASTP hit, evidence for *Mesodinium*’s genetic code, and an alien-index score detecting affinity for cryptophytes in our database^150^ were tabulated (Data S1H). We selected genes with an alien index greater than 10, yielding 136 putative HGT candidates. BLASTP hits for each gene were manually inspected, and genes with strong hits to cryptophytes were further inspected to ensure they were not typically present in ciliates, were present in both the macronuclear and micronuclear assemblies, and displayed evidence of translation in *Mesodinium*’s genetic code.

Genes that met these criteria were further inspected for biological interpretability. Of these, only gene g4460 had a clear interpretation. The amino acid sequence of g4460 was BLASTed against the database again, and all hits with an e-value > 1e-15 were selected. We also manually searched for homologous sequences in the transcriptomes of *Mesodinium chameleon* (8a_k127_2833.p1) and the polar *M. rubrum* species (mrp5_k127_6042.p1) using TBLASTN. These sequences were aligned with MAFFT (FFT-NS-2)^151^ and trimmed with TrimAl^152^ (setting --gappyout). A maximum-likelihood phylogeny was inferred with IQTree2^153^ using the LG+F+I+R6 model with 1,000 bootstrap replicates. We did a Bayesian phylogenetic analysis of gene g4460 using RevBayes^154^ with an LG+I+G6 model. The Markov chain Monte Carlo sampler was run in replicate for 15,000 generations, with 5,000 as burn-in and sampling every 25 iterations. Convergence in each of the marginal posterior distributions was assessed using Tracer 1.7^155^.

#### Searches for tRNAs, tRNA introns, rRNA genes and ITSs

To identify tRNAs and their non-spliceosomal introns we used tRNAscan-SE 2.0 with default settings, for eukaryotic sequences^156^.

To search for rRNA genes and their ITSs we used Infernal 1.1.5^59^ with default parameters.

## Data availability

*Mesodinium rubrum* CBJR05 PacBio HiFi libraries (NCBI SRA PRJNA1341404) from sorted nuclei: MAC (run accession SRR37260499) and MIC (SRR37260498). *M. rubrum* CBJR05 RNA-Seq libraries (ENA PRJEB66165).

*Teleaulax amphioxeia* GCEP01 draft genome assembly (NCBI SRA PRJNA1080848), BRAKER3 gene predictions (Edmond, https://doi.org/10.17617/3.IF79AS).

*Mesodinium rubrum* CBJR05 genome assemblies, gene predictions, repeat and IES annotations (Edmond, https://doi.org/10.17617/3.REWBDY).

FACS data for representative sorting runs (Edmond, https://doi.org/10.17617/3.VI5Q7M).

*M. rubrum* and *T. amphioxeia* genomic and transcriptomic data can be accessed from the PhycoCosm portal^157^explored and mined with the tools available there (at https://phycocosm.jgi.doe.gov/Mesrub2_1 and https://phycocosm.jgi.doe.gov/Telamp1).

Supplementary Data S1A-M (Edmond, https://doi.org/10.17617/3.B4ABRV).

## Acknowledgements

We thank Insa Hirschberg, Moritz Peters, and Frank Chan for access to and assistance with FACS, Abigail Howell for suggestions on cell lysis protocol, Wang Kai and Martina Kolb for sharing reagents, Aurora Panzera and Christian Feldhaus of the Bio Optics facility, Heike Budde and the MPI Biology Genome Center for sequencing support, Andre Noll for computer system administration, Zhongtang Yu for sharing *Entodinium* gene annotations, Agnes Henschen and Dorothee Koch for provision of natural seawater, and Giulio Petroni for discussion about Tlc-related protein HGT.

The work (proposal: 10.46936/10.25585/60001096) was conducted under the auspices of the Community Science Program conducted by the U.S. Department of Energy Joint Genome Institute (https://ror.org/04xm1d337), a DOE Office of Science User Facility, supported by the Office of Science of the U.S. Department of Energy operated under Contract No. DE-AC02-05CH11231.

## Notes

### Competing Interest Statement

The authors have declared no competing interest.

