## Supplementary Information for "Spliceosome loss in the red tide ciliate *Mesodinium rubrum* presents a symbiotic cul-de-sac"

Supplementary Results	3

SR1. Splice junction predictions from RNA-Seq mappings	3

SR2. Presence of a eukaryotic tRNA splicing endonucleases	3

SR3. Helitrons and retrotransposons are the most abundant mobile elements in *M. rubrum*	3

SR4. Superabundant Regulator of Chromosome Condensation 1 (RCC1) proteins	4

SR5. Observations of birefringence in *M. rubrum*	4

SR6. Estimate of the M. rubrum effective population size	5

Supplementary Discussion	5

SD1. Ampliploidy and extensive fragmentation	5

SD2. Potential role of a putative ATP/ADP transporter acquired from Alphaproteobacteria	6

SD3. Consideration of tyrosine tRNAs and the evolution of the *Mesodinium* genetic code	8

SD5. Limitations of published gene models for ciliate genomes	10

SD6. Abundance of RCC1 proteins in ciliates and dinoflagellates	11

SD7. Additional possible hurdles for horizontal gene transfers or their establishment in *Mesodinium*	12

SD8. Retention of IESs compared to loss of introns in *Mesodinium*	12

SD9. Role of trophic lifestyle in intron losses	13

SD10. Role of genome reduction in intron losses	14

SD11. Role of population genetics in intron losses	15

SD12. Role of intron lengths and alternative genetic codes in intron losses	17

References	19

Supplementary Tables	26

Table S1. Genome assembly properties.	26

Table S2. RepeatModeler annotation summary.	27

Table S3. Proteins with the most frequently occurring domains.	28

Table S4. Summary of BUSCO search results.	29

Table S5. Minimum E-values for Infernal snRNA searches.	30

Table S6. Antibodies used for differential immunofluorescence labeling of nuclei.	31

Table S7. Data sources underlying Table 1.	32

Supplementary Figures	33

Figure S1. Blob plots for *Mesodinium rubrum* MIC and MAC assemblies.	33

Figure S2. Distributions of d_s_ and 4-fold synonymous diversity for the MAC genome assembly	34

Figure S3. Birefringence analysis of dinoflagellate and *Mesodinium rubrum* cells.	35

Figure S4. Properties of introns in ciliate genes.	37

Figure S5. IES retention analyses.	38

### **Supplementary Results**

#### SR1. Splice junction predictions from RNA-Seq mappings

Several datasets showed a pronounced enrichment of splice junctions whose lengths are multiples of three (3n), which were also more likely to lack a canonical splice site motif and to have a low edit distance to their flanking regions (Figure 2, Figure S4). These are consistent with the relative enrichment of 3n indels generally seen in coding sequences associated with heterozygosity or genetic variation in the RNA-Seq samples vs. reference genome^1^, and can be contrasted with the relative depletion of 3n introns found in ciliates^2,3^.

*Teleaulax amphioxeia* sequences in the same RNA-seq libraries as *M. rubrum* had splice junctions consistent with conventional introns (both canonical and non-canonical), so we could exclude technical errors in the RNA-seq that could have caused intron detection failure, particularly carry-through of gDNA during library preparation.

Intron metrics from published genome annotations may differ from empirical predictions due to varying methods used for gene prediction and annotation (Supplementary Discussion SD5).

#### SR2. Presence of a eukaryotic tRNA splicing endonucleases

We found five tRNA splicing endonucleases (TSENs) in the predicted proteins from the *M. rubrum* MAC genome (Data S1L). Two pairs of sequences (264 aa and 224 aa, respectively) are identical within each pair; the remaining sequence is 91.8% identical to the longer of the two pairs. Complete catalytic triads (Y, H, K) characteristic of active TSENSs were present in all these proteins, so each could potentially perform the splicing of *M. rubrum*’s tRNA introns.

#### SR3. Helitrons and retrotransposons are the most abundant mobile elements in M. rubrum

With the caveat that the current state of the *M. rubrum* genome assemblies does not enable computational determination of which mobile elements are germline-specific, it is nevertheless possible to obtain useful information about the nature of the most common mobile elements. RepeatModeller annotations (Table S2) revealed “rolling circles” (MIC: 21,409; MAC: 11,370) and retroelements (MIC: 17,457; MAC: 6,324) correspond to the most abundant mobile elements in the assembled *M. rubrum* genomes. As can be seen from counts of Pfam protein domains in Table S3, the most frequent mobile elements correspond to helitrons: 2510 predicted proteins from the MAC genome had helitron-related Pfam protein domain annotations (one of: PF21530, PF14214, PF20209 or PF05970; “DNA helicase Pif1, 2B domain”, “Helitron helicase-like domain at N-terminus”, “Domain of unknown function (DUF6570)” and “PIF1-like helicase”, respectively), 1459 of which correspond to DUF6570. DUF6570 is the distinguishing domain of one of the two main helitron clades, Helitron-like element 2 (HLE2), encompassing helitron variants previously called “Helitron2” and “Helentron”^4^.

1102 predicted *M. rubrum* MAC-genome-encoded proteins had retrotransposon-associated domains (one of: PF00075, PF00078 or PF03372; “RNase H”, “Reverse transcriptase (RNA-dependent DNA polymerase)” or “Endonuclease/Exonuclease/phosphatase family”; Table Domains) in the *M. rubrum* MAC genome. An abundance of retroelements have been observed in some ciliate genomes^5–8^, and numerous helitrons have been observed in the *Oxytricha trifallax* MIC genome^5^ and in *Paramecium tetraurelia^9^*. Helitrons are capable of capturing non-helitron genes and duplicating them when they replicate^10^, which may have contributed to expansions of the number of genes in *M. rubrum*.

#### SR4. Superabundant Regulator of Chromosome Condensation 1 (RCC1) proteins

Mobile elements in *M. rubrum* correspond to the majority of the most abundant protein domains annotated (Data S1J). As far as we can judge from searches of the scientific literature, the domain found in the second most proteins in the MAC genome (914 proteins) and the most proteins the MIC genome (1234), 'Regulator of chromosome condensation (RCC1) repeat' (Pfam: PF13540), has not been associated with mobile elements. We did not observe a tendency for genes encoding these proteins to be adjacent to other mobile element genes in *M. rubrum*. In the majority of cases the RCC1 proteins only contained the RCC1 domain (e.g., 97.6% of MAC-encoded proteins). For discussion about the possible role of these proteins see Supplementary Discussion SD6.

#### SR5. Observations of birefringence in M. rubrum

We analyzed *M. rubrum* and a dinoflagellate, *Akashiwo sanguinea*, as a positive control, to assess the presence of birefringence within their nuclei. As expected, clear birefringence was observed in the dinoflagellate, coinciding with the presence of the large SYBR-positive dinokaryon. However, we also observed some birefringence associated with what appeared to be the MAC of *M. rubrum*, with both the membrane and an internal structure. Slow axis lines are parallel to the nuclear envelope and follow its outline, indicating anisotropy in this region. Lamina of Hutchinson–Gilford progeria syndrome human nuclei exhibit similar birefringence, which was ascribed to ordered lamin microdomains, whereas normal human nuclei do not exhibit birefringence or such ordering^11^. Inside the nucleus, the birefringent structure, which is encircled in red and is of unknown origin (Supplemental Figure S3), has a uniform slow axis direction.

#### SR6. Estimate of the M. rubrum effective population size

We obtained an estimate of 0.0268 for synonymous sequence diversity (π) at 4-fold synonymous sites of genes within collinear genomic segments (distribution shown in Figure S2C), which is within the range of such diversity in the genus *Paramecium* (0.0058 to 0.1599)^12^. To estimate effective population size (N_e_) from π=4N_e_μ, it is necessary to obtain an estimate of the mutation rate (μ, which varies by orders of magnitude across eukaryotes^13,14^) via long-term experimental evolution experiments and genome sequencing or directly by comparisons of parental and offspring genomes. Extremely low estimates of μ have been obtained for the *Paramecium aurelia* complex (1.94×10^-11^ to 2.44×10^-11^)^15,16^ and *Tetrahymena thermophila* (7.61×10^-12^)^17^ by the former approach. Assuming that the mutation rate in *M. rubrum* is similar, the effective population size would be in the range of 275-880 million.

### **Supplementary Discussion**

#### SD1. Ampliploidy and extensive fragmentation

*Mesodinium rubrum* differs from other ciliates in having less DNA in MACs than MICs, apparently lacking somatic genome amplification (ampliploidy). This may relate to their small cell sizes. In ciliates, larger species generally tend to have larger MACs^18^, so, along with genome duplications and having multiple MACs, amplification is thought to be a factor in having enough gene copies available for transcription in these single-celled organisms. Not only is *M. rubrum* smaller than most ciliates, ca. 30 µm length^19^ vs. 50 µm in *Tetrahymena thermophila*, which has ca. 45× amplification^20^, but much of its cell volume is occupied by the kleptokaryon and plastids, with over half of total mRNA transcribed from the kleptokaryon^21,22^. Furthermore, the lack of introns implies that there is no delay for their removal prior to translation of mRNAs, making protein production more efficient. Given that each *M. rubrum* cell has two MACs, additional amplification may simply be unnecessary. Karyorelict ciliates also lack extensive ampliploidy, but their MACs do have more DNA than their MICs and hence do have some degree of amplification^8^.

The only other ciliates we are aware of that also appear to have little genome amplification are two karyorelict ciliates, *Loxodes magnus* and *Loxodes striatus*^8^. In the *L. magnus* genome we observed long tandem arrays of tRNAs similar to those in *M. rubrum*. *Loxodes* cells may be much larger (up to ca. 1 mm length), but instead of amplifying DNA in their nuclei, they seem to predominantly rely upon production of more nuclei, although the number of nuclei is modest compared to the degree of genome amplification in other ciliates. Thus, there are multiple orthogonal ways (nuclear copies, gene duplication, DNA amplification) that can produce additional expressed DNA sequences.

There is no indication thus far that other litostome ciliates may lack ampliploidy, although, as far as we are aware, there is no quantitative data available on nuclear DNA content and limited morphometric data for other litostomes, while stylized illustrations, e.g., in reference 23, may exaggerate the difference in size between MIC and MAC. At least two other litostomes, *Entodinium* and *Balantidium*, have extensive somatic genome fragmentation and nanochromosomes in the MAC, with average lengths about 3-4 kbp^24,25^. On the other hand, other litostome ciliates from ruminants appear to have much larger DNA molecules^26^. We did not detect nanochromosomes in *Mesodinium*, and their MAC contigs are on average an order of magnitude longer than those of *Entodinium* and *Balantidium*, although we are limited by not knowing the telomere repeat sequence of *Mesodinium*.

#### SD2. Potential role of a putative ATP/ADP transporter acquired from Alphaproteobacteria

Although there is no strong evidence for horizontal transfers from other eukaryotes, kleptoplastidic *Mesodinium* species do appear to possess an ATP/ADP transporter acquired from Alphaproteobacteria. Though bacterial gene transfers to eukaryotes present significant problems, such as lacking suitable promoters for expression, a bacterial genic origin would not present the issue posed by introns in eukaryotic genes. How such a gene is incorporated into its germline genome, but also transmitted through to the somatic genome, also poses a problem for the prevailing theory that ciliate nuclear dualism along with sRNAs and epigenetic mechanisms serves as genome defense against foreign DNA integration (critiqued in reference 27). Along similar lines to our searches for HGT in *M. rubrum*, recently a bacterial horizontal gene transfer was reported in *Paramecium* *bursaria*, a ciliate species that harnesses algal endosymbiont photosynthesis, but no algal gene transfers were reported^28^. While the levels of gene transfers from bacteria to ciliates do not begin to approach “you are what you eat”, it is more likely that such transfers will occur from intracellular microbes (cytobionts) by virtue of their proximity. Supporting this, we reported multiple horizontal gene transfers from *Rickettsia* in the genomes of *Loxodes* *magnus*, including a transposase gene and an approximately 9.6 kb cluster of six predicted genes^8^.

The transfer of a putative ATP/ADP transporter from a lineage of intracellular parasites to a kleptoplastidic ciliate is notable. Intracellular parasites, specifically Chlamydiae, are a recurring character in hypotheses of plastid origins due to their genetic contribution in the deep past. Numerous plastid-associated genes of apparent chlamydial origin, including Npt1/Npt2 ATP/ADP transporters, which mediate nucleotide exchange between host and organelle^29–32^. In intracellular Chlamydiae, Npt1 enables parasitic ATP extraction from the host^33^, and a similar role is known for Tlc1 in Rickettsia^34^. The nature of this ancient genetic transfer is debated. Some hypothesize that a tripartite symbiosis among a eukaryotic host, a *Chlamydia*-related bacterium, and a cyanobacterium was responsible for the formation of a stable plastid endosymbiosis^35,36^. Others note that the gene transfer could have occurred in an existing endosymbiosis, allowing the host to better interact with the symbiont^37^.

Regardless of the nature of the gene transfer in ancient eukaryotes, it is clear that the genes transferred play a key role in modern eukaryotic phototrophs. Furthermore, it can be shown to facilitate the establishment of synthetic endosymbioses between cyanobacteria and yeast. In a recent study, the transfer of these ATP transporters from red algae and glaucophytes to yeast enabled an ATP-dependent endosymbiosis between the yeast and a cyanobacterium^38^. The independent transfer of a homologous gene from Alphaproteobacteria into *Mesodinium* could underlie the efficiency of its kleptoplasty. Such a gene could help facilitate energetic transfer between the host and the stolen plastid. If so, this would represent a striking example of convergent evolution, where a transporter initially enabling intracellular parasitism is repurposed in the reverse to exploit the metabolism of a phototroph. The distribution of this gene across *Mesodinium* species is suggestive of this—being present in the kleptoplastidic species *M. rubrum* and *M. chamaeleon* genomic data, and undetected in *M. pulex*, though the latter absence will need to be reevaluated once the *M. pulex* genomes have been sequenced. As a first step towards resolving the nature of the phosphonucleotide transfers, it will be necessary in the future to determine in which membranes this transporter protein is localized in *M. rubrum* and *M. chamaeleon* cells.

#### SD3. Consideration of tyrosine tRNAs and the evolution of the Mesodinium genetic code

The genus *Mesodinium* has a unique and interesting genetic code, translating UAA and UAG as tyrosine (NCBI translation table 29, https://www.ncbi.nlm.nih.gov/Taxonomy/Utils/wprintgc.cgi)^39,40^. Given that diverse alternative genetic codes appear to have evolved independently multiple times in ciliates^41^, and the fact that tyrosine tRNAs anticodons only require a single point mutation of the first anticodon base (to either C or U) for reassignment of UAA or UAG codons, it is not apparent why this genetic code has evolved just once in nuclear genomes (some mitochondrial genomes in other organisms translate UAA as tyrosine; NCBI codes 14 and 33). Though we do not have an explanation for this observation, it is informative to consider the evolution of the tyrosine tRNAs and, particularly, their anticodon base modifications, which influence their codon pairing preferences.

Tyrosine tRNA introns have been reported and experimentally investigated in *Tetrahymena thermophila^42,43^*. Other tRNA introns were also predicted in the *T. thermophila* MAC genome^42^, but are less certain to be present given their lower prediction scores. Enzymes responsible for tRNA intron splicing, TSENs, can also be found in the proteomes of *Tetrahymena*, *Paramecium*, *Stentor*, and *Euplotes* via searches with the relevant Pfam domain in the UniProt database^44^.

Yeast, *Tetrahymena*, and other eukaryotes have a pseudouridine (ψ) base as the central anticodon base in the mature tyrosine tRNAs (i.e., tRNA^Tyr^_GψA_ for yeast and tRNA^Tyr^_QψA_ for *Tetrahymena*), which requires a tRNA intron for the modification from uridine^43,45^. Anticodon base 35 ψ is required for efficient translational readthrough of UAG stop codons in yeast^46^ and both UAA and UAG stop codons in plants^47^.

Unlike yeast and plants, *Mesodinium rubrum* has directly cognate tRNAs for both UAA and UAG codons (i.e., with anticodons 5’-UUA-3’ and 5’-CUA-3’, respectively), which presumably would translate these reassigned stop codons efficiently, as normal codons rather than readthrough codons, thus rendering pseudouridylation of immature tyrosine tRNAs with 5’-GUA-3’ anticodons redundant. There might still be some remaining beneficial redundancy of translational readthrough of the reassigned stop codons UAA and UAG via the near cognate tyrosine tRNAs (i.e., tRNAs with anticodons with a single non-complementary base) with anticodon 35 ψ. Alternatively, this modification may have another role besides translational readthrough (e.g., in stabilizing the modified tRNAs).

Irrespective of whether the specific base pseudouridylation function is necessary, as long as no tRNA genes cognate to the standard tyrosine codons (UAU and UAC) exist, the intron will need to be spliced out to form a functional tRNA. The putative eukaryotic tyrosine tRNA 35 ψ synthase, Pus7 in yeast, has pseudouridylation roles in other ncRNAs^48^, which would favor the continued existence of the putative ortholog of this enzyme in *Mesodinium* without the need to modify the anticodon base. Thus, it is not surprising that we were able to find an obvious Pus7 ortholog (g6965.t1; best BLASTP matches with this in GenBank are to Pus7 proteins) in the *M. rubrum* MAC genome. In principle, tRNAs with anticodons cognate to UAA and UAG do not require a central pseudouridine base, and thus also not an intron, as in the tRNA^Tyr^_GψA_ precursor, consistent with their absence of introns.

In addition to the central anticodon base modification, *Tetrahymena* tyrosine tRNAs appear to have acquired a first anticodon base modification — queuosine (Q) — i.e., tRNA^Tyr^_QψA_, that leads to avoidance of UAA and UAG codon readthrough, thus avoiding competition with tRNA^Gln^_UUA_ and tRNA^Gln^_CUA_ for the same codons that have been reassigned from stops to glutamine^43^. Since *M. rubrum* translates UAA and UAG as tyrosine, there is no need for the avoidance conferred by the *Tetrahymena* queuosine tyrosine tRNA anticodon base, and thus we would predict that anticodon base 34 is not modified in this manner in *Mesodinium*.

Finally, we note that though *Mesodinium* has multiple copies of both UAA and UAG-cognate intronless tRNAs, any of which could mutate to recognize the standard UAU and UAC codons with just a single base anticodon mutation, we have not observed such an occurrence. Based on ribosome profiling of a Pus7 knockout in yeast it was proposed that anticodon 35 ψ does not lead to better translation of UAU and UAC as tyrosine than 35 U^46^. Thus, it remains an open question as to whether the presence of the tRNA introns, and potential pseudouridylation of anticodon position 35 is necessary at all in *Mesodinium*. This question and the identity of the standard tyrosine tRNA anticodon base 35 in *Mesodinium* would be worth investigating in future with ribosome profiling and other relevant experiments. It may shed more light on the nature of tyrosine translation under the standard genetic code, and why UAA and UAG reassignment to tyrosine, via evolution of tRNAs like those in *M. rubrum*, has not occurred in other organisms despite their readthrough of these codons as tyrosine.

#### SD5. Limitations of published gene models for ciliate genomes

Available gene predictions for ciliate genomes were produced by a variety of methods and investigators.

We considered the gene models and published intron annotations from well-studied laboratory model species with gene-dense MAC genomes, particularly *Paramecium tetraurelia*, *Tetrahymena thermophila*, and *Oxytricha trifallax*, to be the most reliable, because the gene models are supported by multiple evidence modes (e.g., https://doi.org/10.1101/2024.01.31.578305), and have been curated by active communities of users. In the case of *Oxytricha*, the gene-sized nanochromosomes of the MAC genome also aided accurate gene prediction. The remainder have issues that affect their use in comparative analyses.

The *Blepharisma stoltei* and *Loxodes magnus* genomes were assembled and annotated in previous studies from our group. Because of their unusually short introns, which poorly fit available gene models, we annotated splice junctions empirically from RNA-Seq mappings, and then predicted genes from an artificial “intronless” version of the genome. Not all genes (and hence introns) will be represented in the RNA-Seq dataset, so there are possible unanticipated gene prediction artifacts or systematic biases.

*Pseudokeronopsis* spp. The published gene annotations appear largely spurious; visual inspection of RNA-Seq mappings show little overlap between predicted genes and RNA-Seq coverage. The published gene models have an uncharacteristic enrichment of 3n introns, and a 3’-localization bias of introns. In comparison, introns empirically annotated from RNA-Seq data have a narrower length distribution more typical of ciliates, and without the 3n-enrichment.

*Entodinium caudatum*. Assembly is published (GCA_002087855.3), while annotations were obtained by personal communication. The annotated introns have a longer tailed length distribution than those found empirically by read mapping, a sharp lower length cutoff (32 bp) that would have excluded many real introns, and an unusual bimodal 5’- and 3’-localization bias. The length cutoff suggests inappropriate parameterization of the gene prediction model. Our reannotation of the same assembly with BRAKER3 using RNA-Seq mapping as hints produced more realistic intron metrics (Figure S4). The longest contig in the assembly, Genbank accession NBJL03000001.1 (original contig identifier Ento_MGS_1), has no RNA-Seq reads mapped but many predicted genes; BLAST of selected regions and mmseqs taxonomy confirm that this contig is a methanogen sequence. There are other likely contaminant sequences too.

*Balantidium ctenopharyngodoni*. The published annotation (https://doi.org/10.6084/m9.figshare.24439159) has very few introns (75 total, 0.003 per gene), which appears to be an under-prediction based on our reanalysis of the RNA-Seq data.

*Chilodonella uncinata*. No MAC draft genome. MIC raw reads have been published, but not the assemblies, so we cannot evaluate claims made in a previous publication.

*Nyctotherus ovalis*. Only 48 introns have been reported, but the published genome is only partial. There was insufficient public data for us to perform a reanalysis.

*Enterocytozoon bieneusi*. No public RNA-Seq data were found on SRA. Publications that reported lack of introns only performed genome sequencing (https://doi.org/10.1371%2Fjournal.ppat.1000261, https://doi.org/10.1093%2Fgbe%2Fevq022) and used homology- and modeling-based methods for intron detection, so we could not include them in our current analysis. Note also that other *Enterocytozoon* species do have introns: https://doi.org/10.1038/ncomms1082.

Short introns of ciliates are challenging to model. Gene prediction software typically expects the longer introns of typical model organisms like metazoans, yeast, and plants. RNA-Seq mappings are usually used as hints for gene prediction, but the default settings of mappers like Hisat2 often have an excessively short minimum intron length cutoff that excludes real short introns of many ciliates.

#### SD6. Abundance of RCC1 proteins in ciliates and dinoflagellates

An abundance of RCC1 proteins was previously identified in the genome of the dinoflagellate *Breviolum minutum*, with 189 such proteins identified^49^ (both dinoflagellates and ciliates are classified as alveolates^50^). Metazoan RCC1 proteins bind to nucleosomes and, as their name suggests, alter chromatin state in a non-DNA sequence-specific manner^51,52^. For *B. minutum* it has been suggested that RCC1 proteins contribute to the condensation in the liquid-crystal organization of its chromosomes^53^. RCC1 protein domains were amongst the most numerous domains identified in a proteomic analysis of *Oxytricha* MICs, but were not observed in the *Oxytricha* MAC proteome within the limits of detection^54^. Such proteins may thus be involved in chromatin condensation in ciliate germline nuclei. It is also possible that such proteins are associated with mobile elements that are yet to be defined.

In both the *Oxytricha* MAC and MIC proteomes WD40 domains (Pfam PF00400; the fifth most abundant domain in the *M. rubrum* MIC genome; Table S3) are more highly represented than those containing RCC1 domains, and, in the MAC, were observed to be more abundant than histone domains^54^. Both RCC1 domains and WD40 domains fold into 7-bladed beta propellers but are considered different by structural biologists^55^. In *Paramecium* a WD40-domain containing protein (PtCAF1), which is upregulated during development and affects new MAC genome development, is thought to be a histone chaperone^56^. So, the abundance of WD40 and RCC1 domain-containing protein genes in *M. rubrum* may reflect a beneficial role for numerous copies in chromatin organization.

#### SD7. Additional possible hurdles for horizontal gene transfers or their establishment in Mesodinium

The distinctive alternative genetic code of *Mesodinium* (UAR=Tyrosine, instead of Stop) is one potential additional hurdle for gene transfer that would result in overextension of proteins transferred to this clade from standard genetic code organisms. Considering ciliates with algal endosymbionts, *Paramecium bursaria* likewise has an alternative nuclear genetic code with a reassigned stop codon (UAR=Glutamine)^57^ that would lead to the same problem, but this problem does not exist in the standard genetic code *Stentor pyriformis*^58^.

The transcriptionally inactive and densely packed ciliate germline nucleus is a natural, passive barrier to HGT. Non-mobile-element gene transfers to the actively transcribed, more accessible somatic genome will be diluted out across cell divisions and lost when somatic genomes are formed anew from germline genome copies during sex. Epigenetic mechanisms that contribute to DNA excision during somatic genome development in ciliates could also prevent any genes successfully incorporated into the germline from being present in the somatic genome, followed by their subsequent pseudogenization and disappearance.

#### SD8. Retention of IESs compared to loss of introns in Mesodinium

Analogous to introns, which are excised from RNA transcripts, IESs are excised from soma-destined (MAC-destined) genomic DNA and persist in the germline (MIC) genome. This leads to distinct germline and soma genome versions. The germline genome participates in meiotic sex and is inherited by the next sexual generation, but is not used for gene expression. In contrast, the “working”, actively transcribed somatic genome is not inherited by the next sexual generation.

Unlike introns, which cannot be generated by spliceosomes that operate upon RNA rather than DNA, ancestral IESs in DNA likely were generated by the same ancestral transposases that remove them during new MAC genome development (PiggyBac transposases responsible for IES excision in *T. thermophila* and *P. tetraurelia* are thought to have lost their ability to generate new transposons as they are no longer present as genes within transposons, and hence are referred to as “excisases”). The very first IESs may have been the transposons that gave rise to extant domesticated transposases, with the transposon/transposase system cleaning up after themselves to produce transposon-free MAC genomes. Besides IESs being largely hidden from selection in the MIC genome and thus able to accumulate, an additional reason why *M. rubrum* has plenty of IESs but is devoid of introns may be that ancestral *M. rubrum* transposases provided the means to continue generating new IESs, whilst introns, incapable of being produced by the spliceosome, were lost. A caveat for this is that such ancestral transposases would need to have been actively transposing in the ancestral MIC genomes.

An important contributor to IESs in *M. rubrum* may be secondary mobile elements that have exploited existing IES excisases for their own replication. Though not as abundant as in *Blepharisma stoltei* and some *Paramecium* species, we also observed signs of potential IES blooming due to MITE non-autonomous mobile elements (MITIES, IESs that are MITES^7^), i.e., short IESs a few hundred bp in length, with pronounced length peaks due to their rapid proliferation (Figure 4). The total amount of genomic DNA derived from these elements appears to be modest (on the order of megabases). However, ancient blooms may have made more substantial contributions before mutating beyond recognition into the ocean of genomic repeats. It is plausible that the ancestral *Mesodinium* genome in which the spliceosome was lost was substantially smaller prior to subsequent expansions due to MITIES and other mobile element activity.

#### SD9. Role of trophic lifestyle in intron losses

Mainstream perspectives on the reasons for large-scale intron losses may be biased by the initial discoveries of extreme intron losses in pathogenic eukaryotic microbial parasites, particularly those with smaller genomes. With subsequent accumulating genomic evidence, the relationship between genome size and trophic lifestyle within eukaryotes is not clear-cut. Parasites can have very large genomes, e.g., 1 Gbp in the parasitic nematode *Aplectana* *chamaeleonis^59^*; within angiosperms, parasites have larger genomes than non-parasites (mode of 9.89 Gbp vs 0.6 Gbp^60^). Parasitoids are largely free-living predators with parasite-like life history interludes, and thus not expected to have extreme genome reduction, e.g., *Nasonia vitripennis*, a parasitoid wasp, has a 297 Mbp genome, substantially larger than the non-parasitoid model insect *Drosophila melanogaster*^61^.

Assuming that no reversion from parasitism has occurred, parasitism is also not a prerequisite for extensive intron loss, as demonstrated by the intron-poor, free-living alga *Cyanidioschyzon* *merolae*. Under a similar assumption, the genus *Mesodinium* indicates that the *M. rubrum’*s parasitoid lifestyle is not the reason for complete spliceosome and intron loss, since the heterotrophic *M. pulex* also appears to have lost its spliceosome. Endosymbiosis is also not a prerequisite for intron losses, since endosymbiotically-derived organelles like the *Bigelowiella natans* nucleomorph still contain plenty of introns, albeit small ones^62^ (see Table 1), whereas some parasitic eukaryotes, *C. merolae*, and *Mesodinium* have lost most or all introns.

#### SD10. Role of genome reduction in intron losses

Like trophic lifestyle, the association between genome reduction and intron losses is not especially obvious given multiple counterexamples (Table 1): the nucleomorph genome of *Bigelowiella* *natans* has plenty of introns (2.6 per gene)^62^; parasitic eukaryotes with small genomes may be intron rich, e.g., *Plasmodium falciparum* (23.5 Mbp genome^63^, but thousands of introns^64^, and the animal *Intoshia variabili* (15.3 Mbp, >26,000 introns^65^). The genomes of *Saccharomyces cerevisiae* (12 Mbp) and *Schizosaccharomyces pombe* (13 Mbp) are a similar size, yet the number of introns these genomes contain differs by an order of magnitude (315 vs. 5,334 introns). Larger genomes may be intron poor, like *Trichomonas vaginalis* (180 Mbp genome, ~38,000 genes)^66^ but only 63 introns in transcribed genes^67^). Interestingly, some of the *T. vaginalis* introns are very short (23 nt), approaching those of *Paramecium tetraurelia* (25 nt length mode)^3^, though the latter has tens of thousands of introns^68^.

Karyorelict ciliates aside, ciliate somatic (MAC) genome sizes are moderate for eukaryotic genomes (~20-100 Mbp). Their germline (MIC) genomes are much larger, containing on the order of 10 Mbp to 1 Gbp of additional sequence, depending on the species, with the amount of DNA comprising mobile elements and repetitive DNA varying greatly^5–7,20,69^. For the technical reasons already explained, it is not possible to provide an exact estimate of the *M. rubrum* MAC genome size based on the data presented, but it is likely to be on the order of 100 Mbp, which is considerably larger than the genomes of most other intronless or intron-poor eukaryotes (but similar to *Trichomonas* genomes).

Substantial MAC genome size expansions have occurred in at least two ciliate lineages. The genus *Paramecium* has a complex history of whole genome duplications^6,70^, with up to three in *Paramecium tetraurelia* (72 Mbp MAC genome). Recently, it was proposed that a whole genome duplication occurred in the lineage leading to the heterotrich *Stentor coeruleus^71^*.

Sequenced genomes are a snapshot of evolutionary history and genome sizes are dynamic, so intron losses could have occurred in smaller ancestral genomes, prior to their subsequent enlargement. Though the karyorelict ciliate *Loxodes* *magnus* possesses a large MAC genome (706 Mb) it has tiny introns (length mode 17 nt; Table 1), but it is plausible that the common ancestor of the sister karyorelict and heterotrich clades possessed a reduced MAC genome with tiny introns. That ancestral genome could have resembled that of the free-living heterotrich *Fabrea salina*, the smallest published ciliate MAC genome (18.4 Mbp), which is roughly twice as gene dense as that of *Plasmodium falciparum* (23.5 Mbp)^72^ and which shares the predominantly 15/16 nt introns and tiny 3’ UTRs characteristic of heterotrichs^40,73^. The MIC-specific portion of the common ancestral karyorelict/heterotrich MIC genome may not have been subject to the selective constraints that led to MAC genome size reduction, and thus junk DNA accrued. Subsequently, in losing the capacity for MAC nuclear division during cell replication, all such junk DNA was included in the large *L. magnus* MAC genome^8^.

It is also possible that some MAC genome size expansion has occurred in *M. rubrum* via proliferation of largely benign mobile elements, like the self-inactivating retroelements in the *Blepharisma stoltei* MAC genome^7^. Both retroelements and helitrons are abundant in *Mesodinium* *rubrum*, but it will be necessary to obtain very clean MAC and MIC separations to ascertain whether and to what degree they are present in the MAC genome.

#### SD11. Role of population genetics in intron losses

Despite the analogy between spliceosomal introns and IESs, and both being present in the germline genome of most ciliates, the former are apparently absent and the latter are abundant in *M. rubrum*. Furthermore, *M. rubrum* also possesses non-spliceosomal tRNA introns and ribosomal RNA internal transcribed sequences. This requires some consideration under the prevailing idea that genome size and associated features like the content of repeats and other non-coding DNA are governed by the same fundamental evolutionary forces^74^.

In a population genetic framework^69,70^, assuming no possibility of further intron gains, the rate of loss would be the product of the rate of introduction of loss-of-intron alleles times the probability of fixation of such an allele. The rate of production of such alleles, assuming diploidy, is 2Nμ_L_, where N is the population size and μ_L_ is the intron loss mutation rate. The probability of fixation of the loss allele is p_fix_ = [1 – exp(-2sN_e_p)]/[1 – exp(-2sN_e_)], where N_e_ is the effective population size, p = 1/2N is the initial frequency of the mutant allele (lost intron), and the selection coefficient, s, can be viewed as the mutational/energetic advantage of such an allele which is equal to the product of the per-nucleotide site mutation rate and the number of sites whose proper identity is necessary for proper identification of both splices sites and branchpoint by the spliceosome (on the order of 10) plus some unknown but small term related to the cost of carrying and transcribing an intron.

Drawing from such theory, one can also consider the equilibrium frequency of introns per nucleotide site (which could be called the occupancy, the average number of introns per coding site) that would result from a balance between the input via intron-gain alleles by mutation and fixation and the output due to loss mutation and fixation. The equilibrium frequency is 1 minus the equilibrium frequency of absentees, where the latter is (μ_L_/μ_G_) × exp(2N_e_s) / [1 + (μ_L_/μ_G_) × exp(2N_e_s)]. In the absence of selection, the equilibrium frequency of introns per site is μ_G_ / (μ_G_ + μ_L_), the neutral expectation. The ratio of μ values is the net pressure due to mutation (equal to 1.0 if there is no bias in gain or loss). The exp() term is the ratio of fixation probabilities, so the product of these two terms is the net mutation–selection pressure in the direction of loss of intron alleles. In summary, the level of intron occupancy within a species is a function of the relative strengths of the physical/mutational forces that result in gain vs. loss alleles, as well as the strength of the downstream selective forces relative to the power of genetic drift.

*M. rubrum* is globally distributed^77^, reported as far apart as both eastern and western US seaboards, as well as the Arctic^78^ and Antarctic oceans^21^. The ancestor of *M. rubrum* could conceivably have had tremendous peak population sizes like blooms of extant *M. rubrum* that span hundreds of kilometers, allowing observation from space (Figure 1B). Presuming that, like extant *M. pulex*, ancestral mesodinia were not specialized upon a single prey species, their effective population sizes may have been even larger than extant *M. rubrum*. The population-genetic explanation would thus be that the large effective population sizes favored elimination of introns in the genus *Mesodinium*. However, *M. rubrum*’s prey species, *Teleaulax amphioxeia*, possessing tens of thousands of introns, also needs consideration (Table 1).

Peak population sizes like the algal blooms in Figure 1B are substantially larger than effective population size, which is the harmonic mean of population sizes across time^79^ and thus strongly dependent upon population bottlenecks. Thus, with a boom-and-bust lifestyle, the ancestral *Teleaulax* effective population size may have been much smaller than that of ancestral *Mesodinium* and continued through till present, leading to many introns. Consistent with the hypothesis that the genomes of cryptophytes in general may be intron-rich, is the fact that another cryptophyte, *Guillardia* *theta*, has numerous introns^80^ (132,894; i.e., 5.3 introns per gene; GenBank accession GCA_000315625.1).

#### SD12. Role of intron lengths and alternative genetic codes in intron losses

Constrained, narrowly selective spliceosomes may favor intron loss; a population genetics model predicts that introns with stricter splice site recognition requirements will be less frequent^76^. Consistent with this, intron-poor eukaryotes such as *C. merolae*, *Giardia* (type A introns) and *Trichomonas* have highly conserved motifs extending beyond the usual GT-AG splice sites^67,81,82^.

Narrow intron length distributions also reflect stricter splicing recognition requirements. The short introns of ciliates have been useful in elucidating general properties of introns influenced by NMD^2,3^. The shortest known introns (15 nt mode) belong to heterotrich ciliates^73,83^ and their sister clade, the karyorelicts (17 nt mode)^8^. Furthermore, aside from *Mesodinium* at zero, these two clades also have the lowest intron densities known among ciliates, e.g., 0.076 and 0.29 introns per gene, respectively, for the karyorelict *Loxodes magnus* and the heterotrich *Blepharisma stoltei*, versus the oligohymenophoreans *Paramecium tetraurelia* (1.6 introns per gene; 25 nt length mode) and *Tetrahymena thermophila* (4.8 introns per gene; 56 nt length mode)^2,73,83^ (Table 1).

Outside ciliates, extremely short introns are also found in nucleomorphs and mikrocytids^84^. The mikrocytids are of particular interest because their tight intron length distribution (predominantly 16, 17 nt) and low intron densities (224 introns in 14,372 genes from *Mikrocytos* *mackini*, i.e., 0.0156 per gene^84^) resemble the heterotrich and karyorelict ciliates. Furthermore, their genomes are not especially small (Table 1), but they appear to have lost most, though not all, spliceosomal proteins.

An additional factor that could have favored intron losses in *Mesodinium* and other ciliates is the use of alternative genetic codes where stops are reassigned to amino acids. The probability of a mutation that “exonizes” the intron via a mutation of the donor, acceptor, or lariat site becomes much higher, since it reduces the probability of introducing a premature stop codon that would lead to protein truncation. In the case of the genus *Mesodinium* there is just a single stop codon instead of the three of the standard genetic code (see Supplementary Discussion SD3). If this genetic code was in use prior to intron losses in this genus, it could have contributed to exonization.

But what happens if the genetic code has context-dependent stops? In such organisms, e.g., the trypanosomatid *Blastocrithidia* nonstop, the heterotrich ciliate *Condylostoma magnum*, and the ciliate class Karyorelictea, the stop codon(s) can also be translated as an amino acid, depending on the surrounding sequence context. This is thought to be governed by the proximity of the ambiguous codon to the 3’-end of the mRNA transcript^40^. If an intron that contains an in-frame context-dependent stop is not spliced out, it would be interpreted as coding if it is close to the 5’ end of the gene, but if it is close to the 3’ end of the gene, it would be interpreted as a stop. Therefore, there would be selection against introns closer to the 5’ end of genes, because nonsense-mediated decay would not be able to effectively deal with the erroneously translated products. This is consistent with *Loxodes*, which uses a “stopless” genetic code, where the intron locations are, unusually, biased towards the 3’ end of genes (Figure S4), unlike other ciliates where it is either uniform or biased towards 5’ end. We speculate that this could function as a “conveyor belt” in organisms with “stopless” genetic codes: selection acts against introns close to the 5’ end of genes, and those that remain at the 3’ end are lost by recombination (RTMIL), leading to an overall net loss of introns.

The argument against this having been the mechanism for intron loss in the *Mesodinium* ancestor is that *Mesodinium* does not have an ambiguous genetic code, and ciliates that do have stopless codes, notably *Loxodes*, still have introns (albeit at low densities), including 3n introns.

### **Supplementary Tables**

#### **Table S1.** Genome assembly properties.

|  | **MAC** | **MIC** |
| --- | --- | --- |
| **Size (Mbp) before GC filtering** | 201 | 356 |
| **Size (Mbp) after GC filtering** | 191 | 347 |
| **Fold coverage** | 63 | 41 |
| **Number of scaffolds** | 9,549 | 9,997 |
| **N50 (kbp)** | 26.6 | 45.5 |
| **Number of genes** | 59,796 | 89,847 |
| **Genes with RNA-seq (≥ 1 AUGUSTUS hints)** | 28,958 | 36,620 |
| **Mean gene lengths (bp)** | 1,294 | 1,023 |

#### **Table S2.** RepeatModeler annotation summary.

|  | **MAC *de novo*** |  | **MIC *de novo*** |  |
| --- | --- | --- | --- | --- |
|  | **Number** | **Length (bp)** | **Number** | **Length (bp)** |
| **Retroelements** | **6,324** | **6,223,451** | **17457** | **11,610,643** |
| **1. SINEs** | 198 | 45,482 | 351 | 82,543 |
| **2. LINEs** | **4,655** | **5,161,494** | **11,753** | **8,315,408** |
| **2.1. L2/CR1/Rex** | 903 | 1,628,578 | 858 | 1,399,817 |
| **2.2. RTE/Bov-B** | 1,991 | 1,622,233 | 4,561 | 2,711,701 |
| **3. LTR elements** | **1,471** | **1,016,475** | **5,353** | **3,212,692** |
| **3.1. Gypsy/DIRS1** | 1,471 | 1,016,475 | 5,353 | 3,212,692 |
| **4. DNA transposons** | **344** | **191,289** | **1,346** | **397,549** |
| **5. hobo-Activator** | 150 | 33,445 | 1,127 | 212,947 |
| **6. Tc1-IS630-Pogo** | 65 | 78,724 | 96 | 107,465 |
| **7. Rolling circles** | 11,370 | 12,136,028 | 21,409 | 14,172,951 |
| **8. Unclassified** | 205,989 | 98,003,059 | 514,611 | 228,113,183 |
| **9. Total interspersed repeats** |  | **104,417,799** |  | **240,121,375** |
| **10. Small RNA** | 859 | 505,856 | 1,293 | 672,495 |
| **11. Satellites** | 125 | 130,696 | 115 | 114,281 |
| **12. Simple repeats** | 39,794 | 3,247,342 | 57,431 | 5,664,398 |
| **13. Low complexity** | 12,603 | 815,531 | 15,046 | 991,273 |

Subtotals are in gray.

#### **Table S3.** Proteins with the most frequently occurring domains.

|  | **MAC** | **MIC** | **Pfam ID** | **Pfam description** |
| --- | --- | --- | --- | --- |
| 1. | 914 | 1234 | PF13540 | Regulator of chromosome condensation (RCC1) repeat |
| 2. | 1459 | 1152 | PF05970 | PIF1-like helicase |
| 3. | 810 | 816 | PF00078 | Reverse transcriptase (RNA-dependent DNA polymerase) |
| 4. | 815 | 815 | PF13358 | DDE superfamily endonuclease |
| 5. | 593 | 730 | PF00400 | WD domain, G-beta repeat |
| 6. | 718 | 726 | PF00075 | RNase H |
| 7. | 801 | 646 | PF14214 | Helitron helicase-like domain at N-terminus |
| 8. | 497 | 496 | PF00069 | Protein kinase domain |
| 9. | 601 | 484 | PF20209 | Domain of unknown function (DUF6570) |
| *10.* | *409* | *425* | *PF13639* | *Ring finger domain* |
| 11. | 543 | 417 | PF21530 | DNA helicase Pif1, 2B domain |
| *12.* | *479* | *383* | *PF00443* | *Ubiquitin carboxyl-terminal hydrolase* |
| 13. | 311 | 263 | PF00226 | DnaJ domain |
| 14. | 233 | 257 | PF13499 | EF-hand domain pair |
| 15. | 170 | 158 | PF03372 | Endonuclease/Exonuclease/phosphatase family |
| Total | 15959 | 18040 |  |  |

Table rows are ordered according to the number of predicted proteins with the given domain in the MIC genome (*de novo* assembly). Colors for mobile element types: green=helitron; blue=retrotransposons; orange=DDE transposons. Italics indicate domains that are either encoded by the same proteins encoding transposases or by proteins whose genes are next to the colored mobile element domains. A notable fraction of genes next to the DDE transposase genes contain Ring finger domains (PF13639). Genes encoding the helitron domain PF14214 also have a substantial fraction of adjacent genes with the Ubiquitin carboxyl-terminal hydrolase (PF00443) domain (comparable numbers to those with the “DNA helicase Pif1, 2B domain” domains) or co-occurring in the same gene. “Total” is the number of proteins with Pfam domains identified, including those not shown in this table. Due to domain copies and sharing within proteins, the total number of proteins in the colored rows is 4,429 and 4,072 for MAC and MIC.

#### **Table S4.** Summary of BUSCO search results.

| **Genome or**  **transcriptome** | **Complete, single-copy or duplicated**  **BUSCOs** | **Fragmented BUSCOs** | **Missing BUSCOs** | **Missing spliceosomal proteins*** | **Total BUSCOs** | **Queried data source** |
| --- | --- | --- | --- | --- | --- | --- |
| *M. rubrum* MAC | 58 | 7 | 34 | 11 | 99 | This study |
| *M. rubrum* MIC | 52 | 10 | 37 | 10 | 99 | This study |
| *M. rubrum* transcriptome | 56 | 4 | 39 | 12 | 99 | Lasek-Nesselquist et al. 2025 |
| *M. chamaeleon* transcriptome | 56 | 3 | 40 | 12 | 99 | Lasek-Nesselquist et al. 2025 |
| *M. pulex* transcriptome | 51 | 3 | 45 | 12 | 99 | Lasek-Nesselquist et al. 2025 |
| *T. thermophila* MAC | 99 | 0 | 0 | 0 | 99 | ciliates.org |
| *Blepharisma stoltei* MAC | 91 | 7 | 1 | 0 | 99 | Genbank |

* Missing spliceosomal proteins are listed in Data S1A.

#### **Table S5.** Minimum E-values for Infernal snRNA searches.

| **Spliceosomal RNA** | **Rfam ID** | ***M. rubrum* MAC** | ***M. rubum* MIC** | ***T. thermophila MAC*** |
| --- | --- | --- | --- | --- |
| U1 | RF00003 | 2.5×10^-4^ | 2.0×10^-4^ | 5.6×10^-25^ |
| U2 | RF00004 | 5.6×10^-4^ | 1.2×10^-3^ | 4.8×10^-33^ |
| U4 | RF00015 | 2.5×10^-5^ | 2.7×10^-5^ | 1.5×10^-20^ |
| U5 | RF00020 | 7.7×10^-4^ | 9.5×10^-4^ | 3.9×10^-17^ |
| U6 | RF00026 | 8.3×10^-5^ | 1.1×10^-4^ | 5×10^-25^ |

#### **Table S6**. Antibodies used for differential immunofluorescence labeling of nuclei.

| **Antibody** | **Type** | **Manufacturer** | **Catalog no.** | **Dilution used** |
| --- | --- | --- | --- | --- |
| Rabbit anti-total histone H3 | Primary | Abcam | ab1791 | 1:100 |
| Rabbit anti-H3K9ac | Primary | Merck | 06-942 | 1:100 |
| Rat anti-alpha tubulin | Primary | Abcam | ab6161 | 1:100 |
| Goat anti-rabbit Alexa Fluor 488 conjugate | Secondary | Abcam | ab150077 | 1:200 |
| Goat anti-rabbit Alexa Fluor 568 conjugate | Secondary | Life Technologies | A11011 | 1:200 |

#### **Table S7**. Data sources underlying Table 1.

|  | **Species** | **Publication** | **GenBank Accession** |
| --- | --- | --- | --- |
| 1 | *Mesodinium rubrum* (host) | This study and https://phycocosm.jgi.doe.gov/Mesrub2_1 |  |
| 2 | *Teleaulax amphioxeia* (prey) | This study and https://phycocosm.jgi.doe.gov/Telamp1 |  |
| 3 | *Blepharisma stoltei* | https://doi.org/10.1073/pnas.2213887120 | GCA_965603825.1 |
| 4 | *Loxodes magnus* | https://doi.org/10.1073/pnas.2400503121 | GCA_946699065.2 |
| 5 | *Paramecium tetraurelia* | https://doi.org/10.1038/nature05230 | GCA_000165425.1 |
| 6 | *Tetrahymena thermophila* | https://doi.org/10.1371/journal.pbio.0040286 | GCA_000189635.1 |
| 7 | *Plasmodium falciparum* | https://doi.org/10.1038/nature01097 | GCA_000002765.3 |
| 8 | *Intoshia variabili* | https://doi.org/10.1016/j.cub.2020.01.061 | (no assembly in Genbank) |
| 9 | *Saccharomyces cerevisiae* | For the numerous associated publications listed in GenBank see: https://www.ncbi.nlm.nih.gov/datasets/genome/GCA_000146045.2 | GCA_000146045.2 |
| 10 | *Schizosaccharomyces pombe* | https://doi.org/10.1093/genetics/iyae007 | https://www.pombase.org/monthly_releases/2025/pombase-2025-11-01/ |
| 11 | *Cyanidioschyzon merolae* | https://doi.org/10.1186/1741-7007-5-28 | GCA_000091205.1 |
| 12 | *Giardia duodenalis* | https://doi.org/10.1099/mgen.0.001117 | GCF_000002435.2 |
| 13 | *Encephalitozoon cuniculi* | https://doi.org/10.1186/1471-2164-14-207 | AEWQ00000000.1 |
| 14 | *Pseudoloma neurophilia* | https://doi.org/10.1016/j.cub.2023.10.034 | GCA_001432165.1 |
| 15 | *Enterocytozoon bieneusi* | https://doi.org/10.1371/journal.ppat.1000261 | GCA_000209485.1 |
| 16 | *Trichomonas vaginalis* | https://doi.org/10.1371/journal.ppat.1013282 | GCA_026262505.1 |
| 17 | *Mikrocytos mackini* | https://doi.org/10.1186/s12915-023-01635-w | GCA_030180075.1 |
| 18 | *Bigelowiella natans nucleomorph* | https://doi.org/10.1073/pnas.0600707103 | DQ158856.1, DQ158857.1, DQ158858.1 |
| 19 | *Guillardia theta nucleomorph* | https://doi.org/10.1038/nature11681 | ADNK00000000.1 |
| 20 | *Hemiselmis andersenii* nucleomorph | https://doi.org/10.1073/pnas.0707419104 | GCA_000018645.1 |

### **Supplementary Figures**

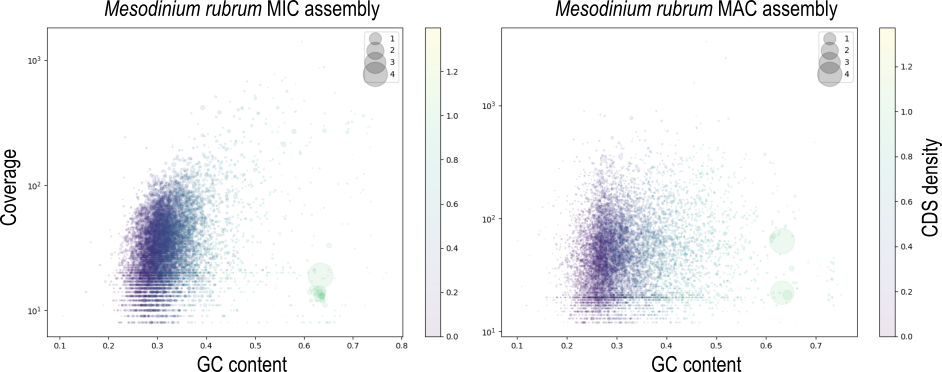

#### **Figure S1**. Blob plots for Mesodinium rubrum MIC and MAC assemblies.

Circle areas are proportional to contig size (legend scale is Mbp).

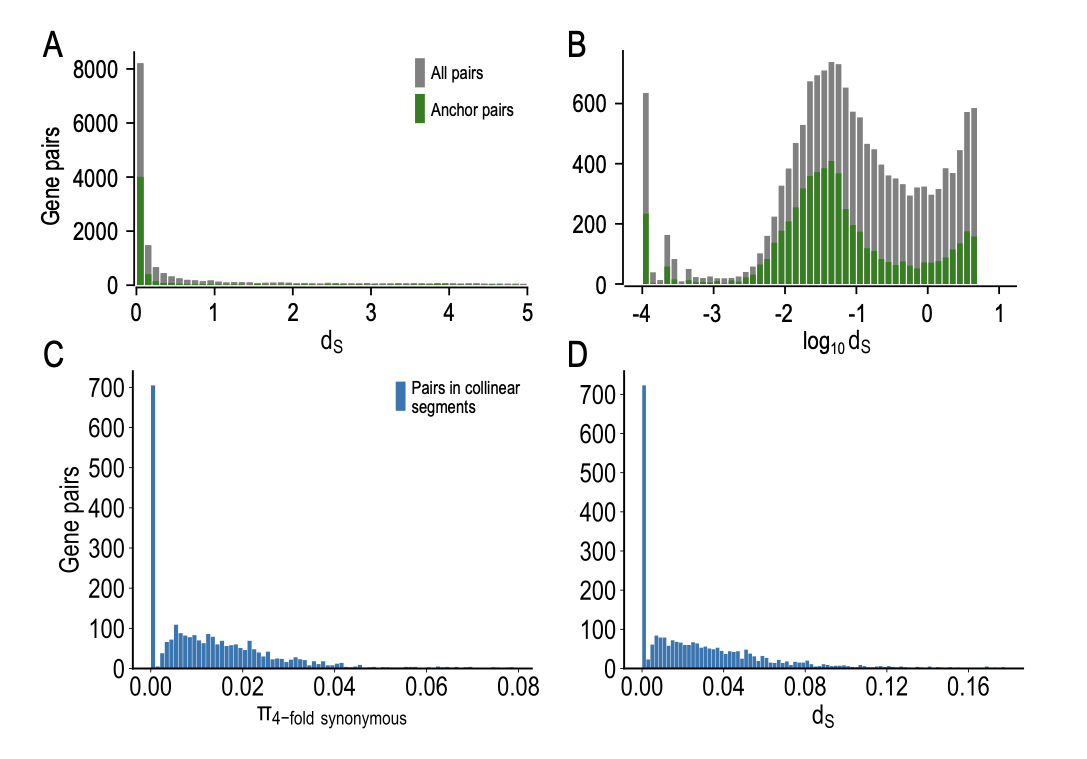

#### **Figure S2**. Distributions of d_s_ and 4-fold synonymous diversity for the MAC genome assembly.

(**A**, **B**) Since the *M. rubrum* genome is not haploid, nor was an inbred strain used, “Gene pairs” includes both duplicated genes (paralogs) and diploid allelic gene variants. Green bars correspond to gene anchor pairs as identified by i-ADHoRe 3 called by the “wgd syn” command. (**C**, **D**) π_4-fold synonymous_ and d_S_ values for just those gene pairs identified as being within collinear segments by wgd v3 via i-ADHoRe 3, i.e., dominated by allelic variants.

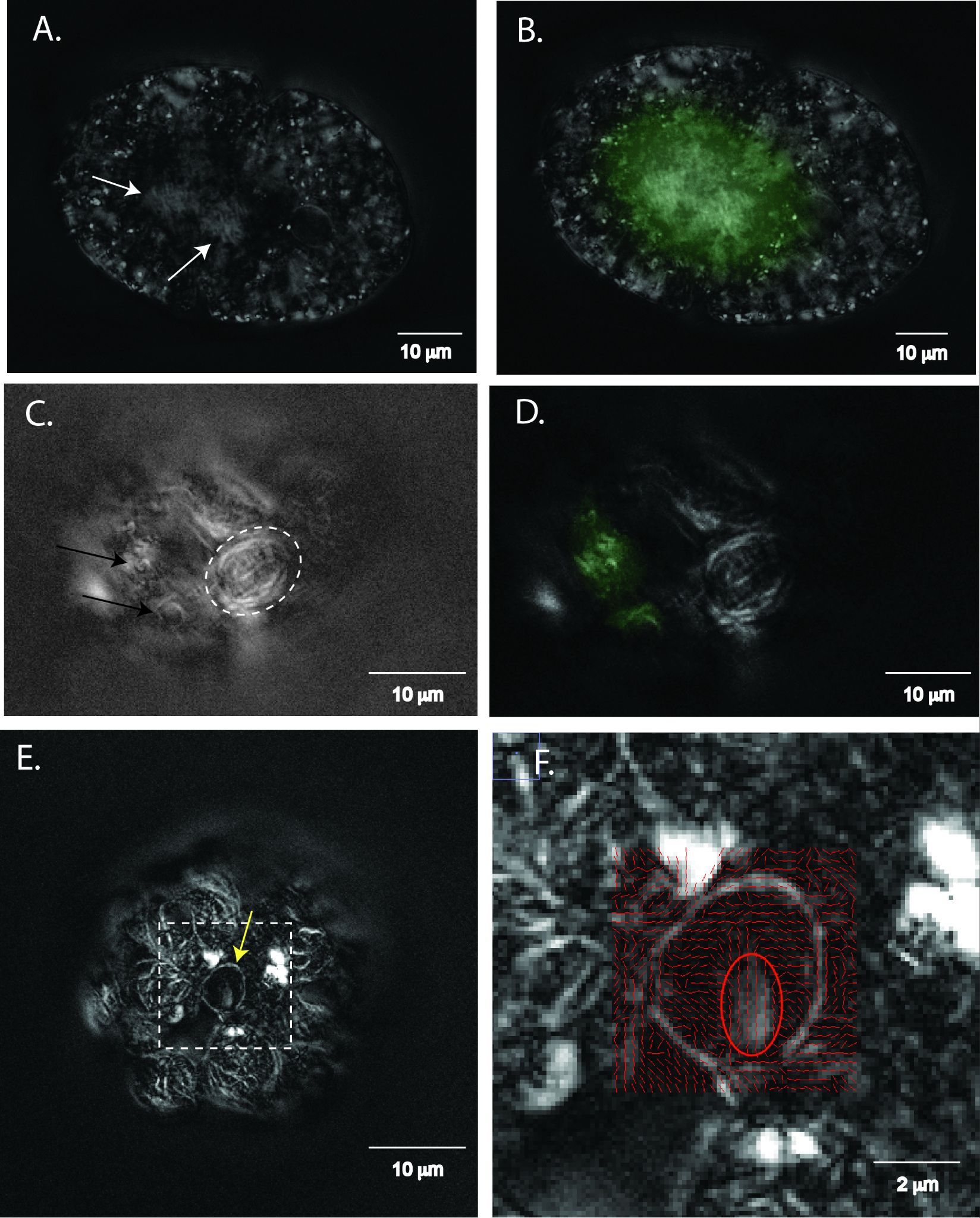

#### **Figure S3.** Birefringence analysis of dinoflagellate and Mesodinium rubrum cells.

Images created by an LC-PolScope (Mehta et al., 2013); birefringent structures cause retardance (differential phase shift) of polarized light, shown here as brightness (grayscale). **A.** An Akashiwo sanguinea (a dinoflagellate, for comparison) cell with numerous internal birefringent structures, including chromosomes within the dinokaryon (white arrows). **B.** Same as in A, with the dinokaryon stained with SYBR Green (green fluorescence). **C.** An M. rubrum cell showing numerous birefringent structures associated with chloroplasts (e.g., white dashed oval) and the ciliate nuclei (black arrows). **D.** Same as in C, showing SYBR Green-stained nuclei (green) correlating with birefringent structures. **E.** An M. rubrum cell with birefringent structures associated with chloroplasts and a macronucleus, revealing birefringence associated with the nuclear membrane (yellow arrow), and an internal birefringent structure. **F.** A close-up of the boxed region from E, showing azimuth (angle of slow axis) lines in red and retardance in gray.

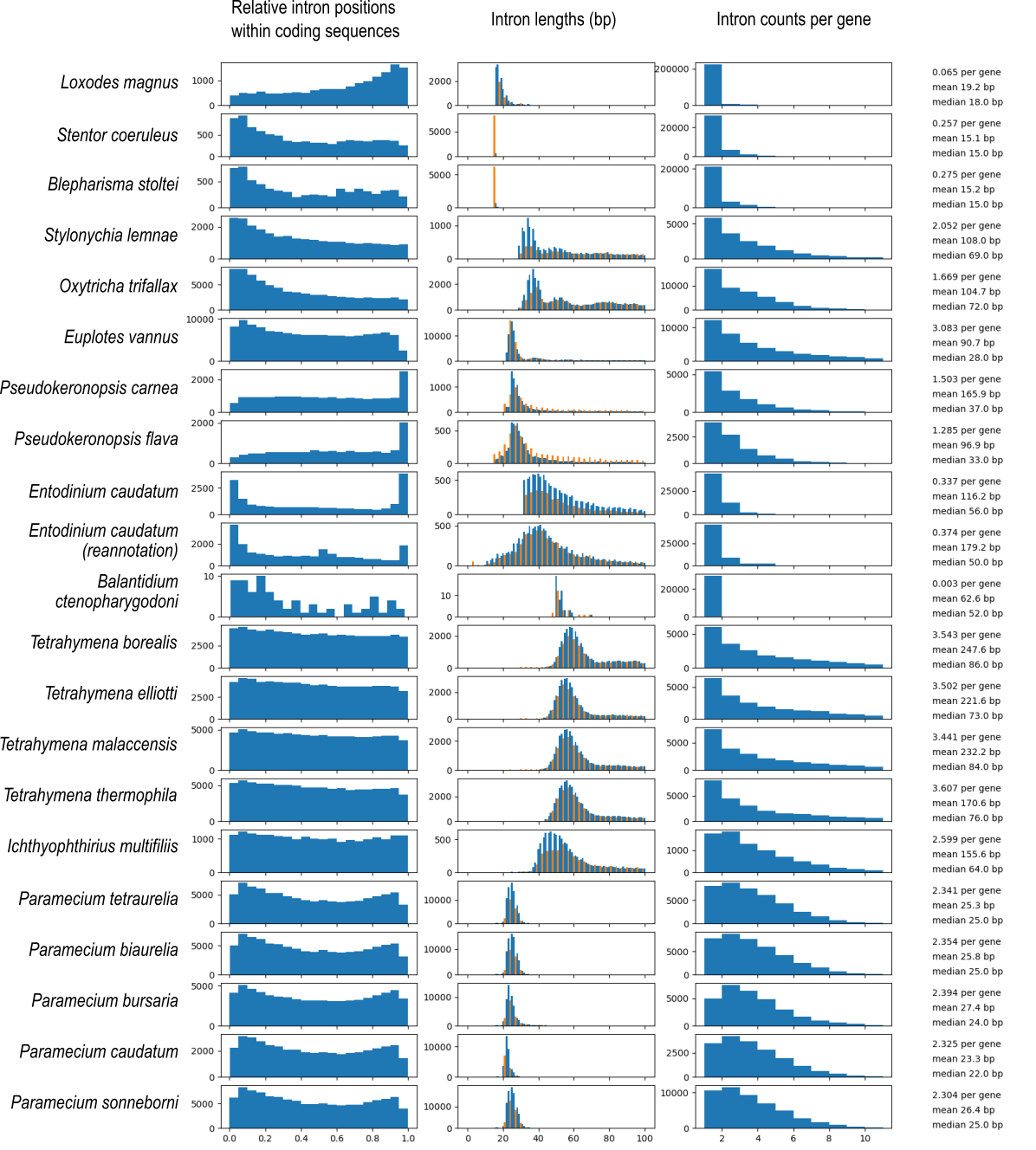

#### **Figure S4.** Properties of introns in ciliate genes.

Histograms of intron position within coding sequences (left), length (middle, up to 100 bp), and per-gene density (right) in published ciliate gene models. Orange: 3n intron length.

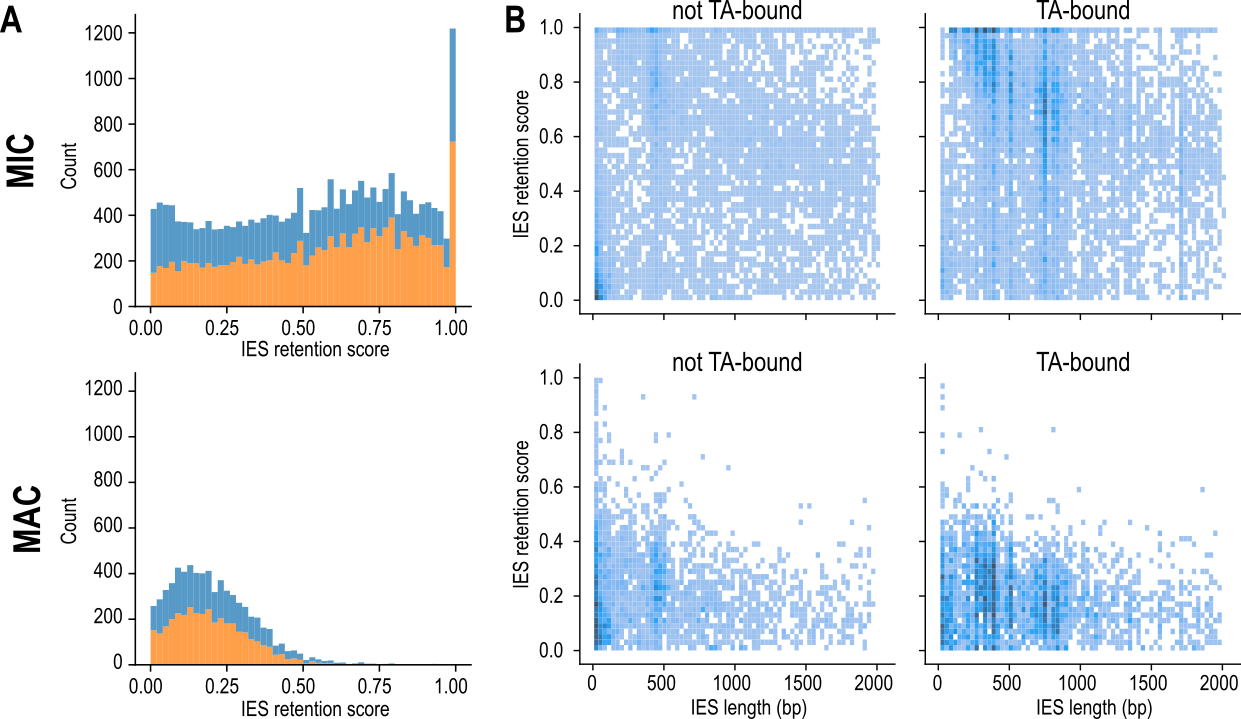

#### **Figure S5.** IES retention analyses.

(A) Histogram of IES retention scores from MIC (above) and MAC libraries (below); orange – TA-bound, blue – not TA-bound. (B) Heatmaps of IES retention scores vs. IES length for MIC (above) and MAC libraries (below).
